# Molecular and physiological characterization of tillering and shade tolerance of dwarf mutants of perennial ryegrass

**DOI:** 10.1101/2024.08.18.606542

**Authors:** Rahul Kumar, Huseyin Yer, Wei Li, Xiangning Jiang, Ying Gai, Hui Duan, Yi Li

## Abstract

Tillering and shade tolerance are important traits in turfgrass, influenced by environmental factors, nutrients, and hormones. Shade stress negatively affects tillering. In this study, two dwarf mutants, *shadow-1* and *shadow-2*, developed via Gamma-ray and fast-neutron mutagenesis, respectively, showed significantly higher tillering than the wild-type under greenhouse conditions. Both mutants demonstrated shade tolerance in plant height, grass quality, and color under 85% and 95% shade conditions, while shade-induced inhibition of tillering was observed in both the mutants and the wild-type. In comparison to wild-type plants under 95% shade conditions, we observed that the cytokinin biosynthetic gene *IPT8* is upregulated, while the cytokinin inactivating gene *CKX2* is downregulated in *shadow-1*. Similarly, the GA biosynthetic genes *CPS1, GA2ox3, and GA20ox1* are upregulated, while the GA inactivating gene *GA20ox8* is downregulated in the *shadow-1* mutant. Furthermore, the ethylene biosynthetic genes *ACS* and *ACO* are also downregulated in the *shadow-1* mutant. Consistently, we observed that wild-type plants exhibit increased GA and reduced CK levels, while *shadow-1* mutant plants have reduced GA but increased CK levels. This explains the *shadow-1* mutant’s shade tolerance in terms of plant height, grass quality, and color. Conversely, the tillering inhibitor genes *CRY1, MAX2,* and *SnRK1* are upregulated in both wild-type and *shadow-1* mutant plants. Our results provide novel insights into the mechanisms behind tillering and shade tolerance in turfgrasses under shade conditions.

## 1. Introduction

Perennial ryegrass (*Lolium perenne* L.) is a highly valued cool-season turfgrass species due to its rapid establishment and attractive appearance, and it is widely cultivated for various purposes such as residential lawns, parks, athletic fields, golf courses, fairways, orchards, and corn/crop fields (Jiang and Huang 2001; Grinberg *et al*. 2016). However, despite its beneficial traits, perennial ryegrass is relatively susceptible to environmental stresses such as drought, salt, and shade (Gardner *et al*. 2002), which can lead to a reduction in turfgrass density and quality (color) (Kebrom and Brutnell, 2007; Zhuang *et al*. 2019). In particular, shade created by trees, bushes, and buildings, is a significant stress for perennial ryegrass (Xing *et al*. 2009). Therefore, developing shade tolerant mutants of perennial ryegrass and understanding the mechanism of the shade tolerance in perennial ryegrass are of great significance for basic and breeding research.

Perennial ryegrass is known to struggle in shade conditions, displaying shade avoidance responses (SAR) such as weak growth, reduced tillering, shoot elongation, and leaf senescence (Franklin and Whitelam, 2005; Li *et al*. 2016; Li *et al*. 2017). Plant hormones play a crucial role in a plant’s response to shade stress (Pierik *et al*. 2004a; Pierik *et al*. 2004b; Das *et al*. 2016; Li *et al*. 2017). Gibberellic acid (GA) is a significant regulator of cell elongation and plant height, and a dwarf phenotype results from impairment of the GA biosynthesis pathway (Hedden 2020; Wu *et al*. 2021; Pearce, 2021; Cheng *et al*. 2022). Increased gibberellin concentration has been reported in beans (Beall *et al*. 1996), sunflowers (Kurepin *et al*. 2007), and *Arabidopsis* under shade stress (Bou-Torrent *et al*. 2014; Liu *et al*. 2016). The expression of GA biosynthesis genes, *GA20ox1, GA20ox2,* and *GA3ox,* is induced by shade stress, leading to increased GA concentration (Hisamatsu *et al*. 2005; Yu *et al*. 2015; Li *et al*. 2017). Shade tolerance is observed in GA-deficient plant mutants (Pierik *et al*. 2004a; Li *et al*. 2017). The *shadow-1* mutant has been used to study the mechanisms of dwarfism and shade tolerance, demonstrating the role of GA hormone in these phenotypes (Li *et al*. 2017). Ethylene (ET) plays a negative role in shade tolerance in plants, while ET-insensitive tobacco shows increased shade tolerance with reduced ET levels (Pierik *et al*. 2004b). Cytokinins (CKs) have a positive effect on shade tolerance in plants, and over production of CKs increases shade tolerance (Gan *et al*. 1995; Robson *et al*. 2004; Xing *et al*. 2009; Xu *et al*. 2010; Schafer *et al*. 2015; Panigrahy *et al*. 2019). The overexpression of the isopentyl transferase (*IPT*) transgene in creeping bentgrass (Agrostis stolonifera) and ryegrass increases endogenous CK content and shade tolerance (Xu *et al*. 2010; Robson *et al*. 2004).

High tillering is a desirable trait in turfgrasses because it is linked to their resistance to wear, damage, and compaction (Carrow *et al*. 1980; Parr *et al*. 1981; Shildrick and Peel, 1984; Lush *et al*. 1990; Mana *et al*. 2021). This characteristic is particularly important for golf greens, where the turf must support the weight of the ball and provide a smooth, even surface (Lush *et al*. 1990). Tillering is controlled by complex interactions between hormones, nutrients, and environmental factors (Ongaro *et al*. 2008; Müller and Leyser, 2011; Leduc *et al*. 2014; Rameau *et al*. 2014; Barbier *et al*. 2015b; Roman *et al*. 2016; Fichtner *et al*. 2017; Le Moigne *et al*. 2018). Auxin, strigolactones (SLs), CKs, and abscisic acid (ABA) are the main plant hormones involved in the regulation of tillering in plants (Ferguson & Beveridge, 2009; McSteen *et al*. 2009; Zhuang *et al*. 2019; Yan *et al*. 2020). Auxin, produced in young tissues, restricts axillary bud development through basipetal transport and may interact with SLs, ABA, and CKs to regulate tillering (Sachs and Thimann, 1967; Cline, 1994; Ljung *et al*. 2001; Blakeslee *et al*. 2005; Yan *et al*. 2020; Shang *et al*. 2021). While auxin up-regulates SLs and ABA biosynthesis genes to inhibit tillering, it acts antagonistically with CKs, which are synthesized in the root and shoot and promote tillering when transported to axillary buds (Kudo *et al*. 2010; Müller and Leyser, 2011; Brewer *et al*. 2009; Yan *et al*. 2020; Shang *et al*. 2021, Tang *et al*. 2023).

Isopentenyltransferases (*IPTs*) control a rate-limiting step in CKs biosynthesis, while cytokinin oxidase/dehydrogenase (*CKX*) controls the endogenous CKs level by degrading CKs in plants (Miyawaki *et al*. 2004; Nordstrom *et al*. 2004; Tanaka *et al*. 2006; Idan *et al*. 2022). *CKXs* genes have been identified in various plant species such as *Arabidopsis* (Werner *et al*. 2003; Werner *et al*. 2006), rice (Ashikari *et al*. 2005), fox millet (Wang *et al*. 2014), and maize (Gu *et al*. 2010). *CKX2* has been reported to play an important role in tiller number through negatively regulating CKs content in plants (Yeh *et al*. 2015; Zhuang *et al*. 2019). Silencing of *CKX2* gene in rice and barley leads to the enhanced endogenous CKs level in plants (Zalewski *et al*. 2010; Yeh *et al*. 2015). Higher endogenous CKs level increases tiller number in plants (Zalewski *et al*. 2010; Yeh *et al*. 2015; Wang *et al*. 2018; Liu *et al*. 2019). *MAX2* encodes an F-box protein required for SLs-mediated inhibition of tillering (Stirnberg *et al*. 2002; Stirnberg *et al*. 2007). A loss of function mutations in the *MAX2* gene produces a high-branching/high-tillering phenotype in *Arabidopsis*, peas, and rice (Stirnberg *et al*. 2002; Ishikawa *et al*. 2005; Johnson *et al*. 2006).

Tillering is one of the turfgrass characteristics that is negatively affected by shade (Kebrom and Brutnell, 2007). Shade decreases tiller number and increases leaf length in the perennial ryegrass (Bahmani *et al*. 2000; Franklin and Whitelam, 2005). Sugar availability in the plant is negatively affected by the shade due to a reduction in photosynthesis (Lunn *et al*. 2006; Yadav *et al*. 2014). Trehalose 6 phosphate (Tre6P) functions as a signal of sucrose availability in plants (Lunn *et al*. 2006; Yadav *et al*. 2014). Tre6P is synthesized by Tre6P synthase gene (*TPS*) and dephosphorylated to trehalose by Tre6P phosphatase gene (*TPP*) (Ganie *et al*. 2019). Experimental evidence suggests that *Tre6P* positively regulates tillering via suppression of sucrose non-fermenting kinase 1 (*SnRK1*) (Simon *et al*. 2018; Baena *et al*. 2020). Sugar signals regulate bud outgrowth through the *SnRK1* gene (Tsai and Gazzarrini, 2014). *SnRK1* is a sensor of energy availability that negatively regulates tillering depending on an energy level to maintain energy homeostasis (Tsai and Gazzarrini, 2014). Sugar availability represses the expression of the *MAX2* gene in plants. Shade, which decreases sugar level, up-regulates *MAX2* resulting in inhibition of buds’ outgrowth (Kebrom *et al*. 2010). Cryptochrome is a photoreceptor that receives shade signals in plants (Zhai *et al*. 2020; Lyu *et al*. 2021). Cryptochrome 1 (*CRY1*) inhibits the shoot branching in the *Arabidopsis,* suggesting a negative role in tillering (Zhai *et al*. 2020; Lyu *et al*. 2021).

Two dwarf mutants, *shadow-1* and *shadow-2*, developed by using Gamma-ray and fast-neutron mutagenesis, were used in this study. Both dwarf mutants exhibited more tillering under light conditions compared to wild-type. In addition, both mutants showed more shade tolerance under 95% and 85% shade conditions compared to wild-type. This study aims to investigate the hormonal and transcriptomic changes responsible for this enhanced shade tolerance and high tillering in the mutants by conducting a combined hormonal and transcriptome analysis. The results of this study provide valuable information on the mechanisms underlying shade tolerance and high tillering in perennial ryegrass, which could have implications for improving the density and quality of turfgrass in shaded areas.

## 2. Material and Methods

### 2.1. Plant materials and mutagenesis

In this study, two high-tillering shade-tolerant dwarf mutants, *shadow-1* and *shadow-2*, were utilized. *shadow-1* was created in our laboratory through gamma-ray mutagenesis of Fiesta-4 seeds, as described in previous studies (Li *et al*. 2016; Li *et al*. 2017). *shadow-2* was developed through fast-neutron (FN) mutagenesis of Fiesta-4 seeds. Wet seeds of ‘Fiesta-4’ perennial ryegrass were subjected to irradiation with 1.0 kr of FN. The irradiated M_0_ seeds were air-dried for 12 hours, sown on the farm, and M_1_ seeds were collected the following year. A total of 300,000 M_1_ seeds were grown in trays containing Promix potting soil. After three weeks of germination, putative dwarf mutants were selected at the three-leaf stage and grown in the greenhouse to confirm their dwarfism and evaluate other phenotypic traits.

### 2.2. Phenotyping of plants under greenhouse conditions

Wild-type, *shadow-1,* and *shadow-2* mutant plants were vegetatively propagated and grown in 7.6 cm plug trays. Plants were maintained at a 5 cm height for six weeks inside the greenhouse with 14/10 hours day/night light conditions. Five tillers for each wild-type and mutants were propagated in plug trays in 10 replications. After six weeks, tiller number and canopy height were measured for wild-type and mutants.

### 2.3. Shade treatment under greenhouse conditions

Following six weeks of growth in full light conditions, ten plugs of wild-type, *shadow-1*, and *shadow-2* plants were transferred to a greenhouse room where shade treatment was given by growing them under 95% shade conditions. The light intensity was measured using a MQ-100 Quantum Meter, Apogee Instruments, Logan, Utah, USA. The tiller numbers were counted four weeks after the continuous shade treatment. A parallel experiment was conducted under full light conditions to compare the effect of shade on the tiller number.

### 2.4. Evaluation of mutants under the shade treatment in the field

Wild-type, *shadow-1*, and *shadow-2* plants were first propagated in 50-plug trays and allowed to grow for 8 weeks under full light conditions in the greenhouse. After 8 weeks, six plugs of each genotype were transplanted into a densely wooded area of the field, where they were exposed to ∼85% shade conditions. The plants were fertilized and irrigated as needed, and their canopy height was maintained at 5 cm by cutting whenever it reached a height of 7.5 cm. Canopy heights and photographs were taken regularly to observe the growth and development of the plants under shade conditions.

### 2.5. Hormonal quantification

Hormonal quantification was done to compare hormone contents between wild-type, *shadow-1,* and *shadow-2* plants under light and shade conditions. Leaf sampling and hormone quantification were performed as described by Li *et al*. (2016). Leaf samples were collected from the wild-type and mutants for hormonal quantification. Leaf samples from 10 plants were pooled for each biological replicate. Three biological replicates were analyzed for each genotype and treatment. About 200 mg of frozen leaf samples were ground, using a mortar and pestle, to a fine powder in liquid nitrogen. Hormone content analysis was carried out using an ultra-high-performance LC-tandem mass spectrometer (UPLC/MS/MS) (Quattro Premier XE ACQUITY Tandem Quadrupole; Waters, Milford, MA, USA).

### 2.6. Identification of differentially expressed genes from transcriptome data

Gene expression analysis was conducted following the procedure described by Li *et al*. (2017). This transcriptome dataset was used in our lab to understand the molecular basis of dwarfism and shade tolerance in the *shadow-1* mutant (Li *et al*. 2017). Transcriptome sequencing data were deposited in the NCBI SRA database under the accession number SRP102018. Clean reads were obtained by removing adapter sequences, then filtering out reads with over 20% low-Q-value (£20) bases, as well as reads with more than 5% ambiguous “N” bases. The clean reads were then aligned to the perennial ryegrass genome assembled by Byrne *et al*. (2015) with default settings in Tophat2 software (Kim *et al*. 2013). Gene expression levels were calculated as reads per kilobase of transcript per million mapped reads. Tillering and shade-regulating genes were identified in perennial ryegrass by BLASTP against the translated perennial ryegrass reference genome for each gene of interest. BLAST analysis was performed using Geneious prime bioinformatics software (Geneious version 2019.1.3).

### 2.7. RNA extraction and qRT-PCR analysis

Leaf tissues were collected from three plants of each genotype for the experiments. The young leaves for each treatment were collected and immediately frozen in liquid nitrogen. Total plant RNA was extracted with a RNeasy Plant Mini Kit, including an RNase-Free DNase set (Qiagen, Valencia, CA, United States), according to the manufacturer’s protocol. RNA purity and concentration were measured using a NanoDrop 2000 spectrophotometer (Thermo Fisher Scientific, Waltham, MA, United States). An iScript^TM^ cDNA Synthesis Kit (Bio-Rad Laboratories, Richmond, CA, United States) was used to synthesize cDNA, and cDNA products were utilized for qRT-PCR assays with SsoFast^TM^ EvaGreen^®^ Supermix (Bio-Rad, Richmond, CA, United States) on a CFX96^TM^ Real-Time PCR detection system (Bio-Rad, Richmond, CA, United States). Primer sequences for all the genes used for qRT-PCR are given in Table 1. The native glyceraldehyde-3-phosphate dehydrogenase (*LpGAPDH*) gene was used as the internal control (Petersen *et al*. 2006). The data were analyzed in CFX Manager^TM^ software. The expression level of each sample was normalized to the expression level of *LpGAPDH* in the same sample.

**Table 1:**
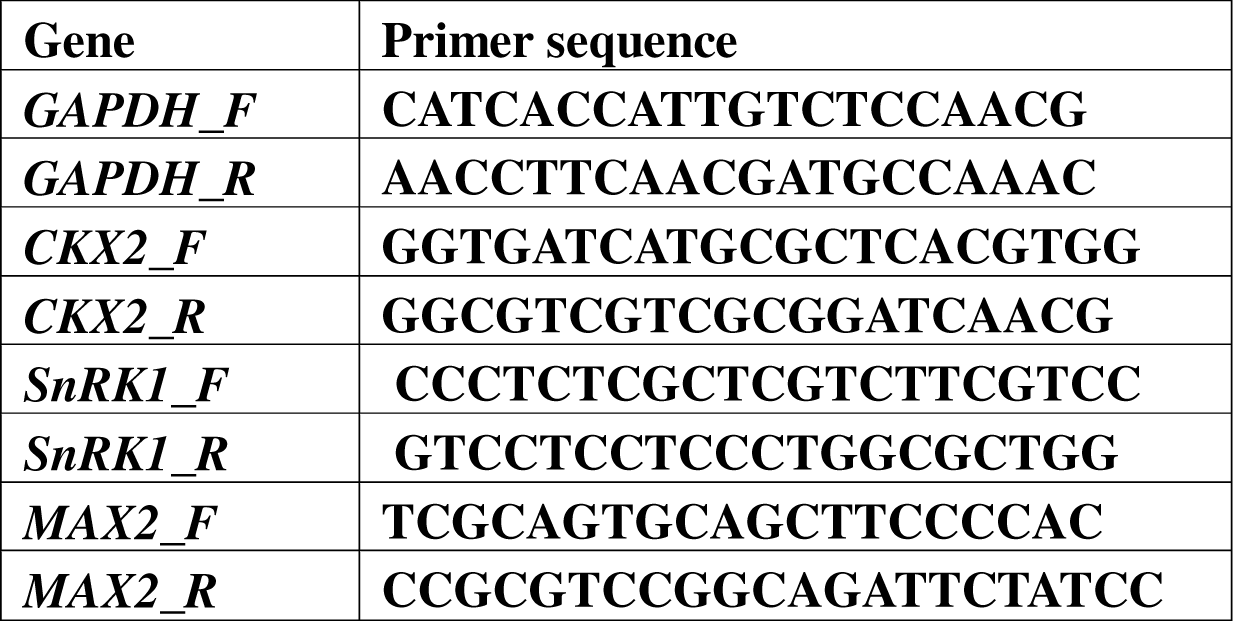
Primer sequences used for qRT-PCR analysis of the selected genes associated with tillering and shade tolerance.

## 3. Results

### 3.1. *shadow-1* and *shadow-2* dwarf mutants had more tiller numbers than wild-type

After six weeks of vegetative propagation, canopy height and tiller number were measured for the phenotypic analysis of wild-type, *shadow-1,* and *shadow-2* mutants under light conditions. The average canopy heights of the wild-type, *shadow-1,* and *shadow-2* mutant plants were 17.3, 9, and 7 cm, respectively (Figure 1A). This result indicated a significant reduction in canopy heights of both mutants compared to the wild-type control. Both mutants had significantly more tillers than wild-type plants. The wild-type plants had an average of 19.1 tillers per plug, while *shadow-1* and *shadow-2* mutants had a significantly higher average of 32 and 29 tillers per plug, respectively (Figure 1B).

**Figure 1.**
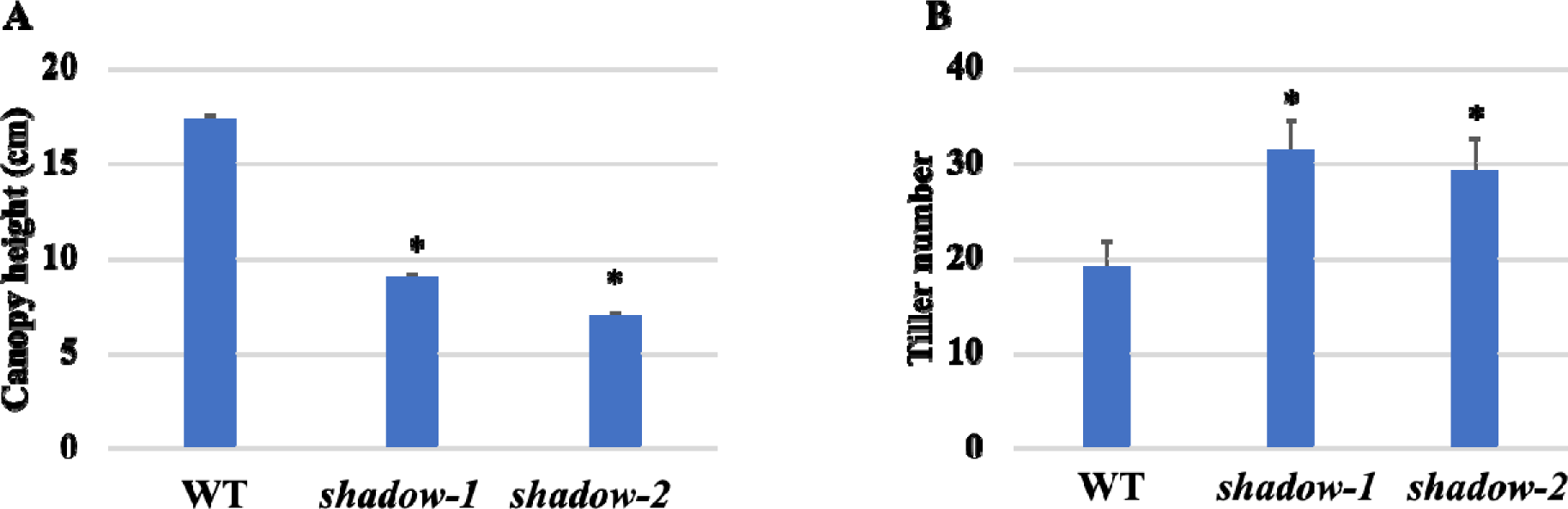
Canopy height and tiller number of *shadow-1* and *shadow-2* mutants under light conditions. Wild-type, *shadow-1,* and *shadow-2* mutants were vegetatively propagated and grown in the greenhouse for six weeks in 7.6 cm plug trays. Five tillers for wild-type, *shadow-1, and shadow-2* mutants were propagated in trays in 10 replications. Canopy height and tiller number were taken as the average of 10 plants. **(A)** Canopy heights of the plants were measured after six weeks of propagation. Both mutants showed a significant reduction in canopy height compared to wild-type. **(B)** Tiller numbers of the plants were measured after six weeks of propagation. All the visible tillers were counted. Both mutants had a significantly higher tiller number compared to wild-type. *Mean differing from wild-type at P < 0.001.

### 3.2. *shadow-1* and *shadow-2* dwarf mutants exhibited shade tolerance, but tillering inhibited in wild-type and mutants under shade conditions

To evaluate the tillering and shade tolerance of *shadow-1* and *shadow-2* mutants, the plants were exposed to 85% shade environments in a densely wooded area (Figure 2B). After transferring the plants to the field, they were maintained at a 5 cm canopy height and were cut whenever they reached a height of 7.5 cm. Over a 2-month period under 85% shade conditions, the wild-type plants started to deteriorate and lost leaf color, while *shadow-1* and *shadow-2* plants remained healthy. Both mutants maintained more tillers than the wild-type plants under 85% shade conditions (Figure 2B). The leaf elongation rates of *shadow-1* and *shadow-2* were significantly lower than wild-type. After 30 months, the wild-type plants had completely died, while *shadow-1* and *shadow-2* plants had maintained most of their tillers (Figure 2B). These results suggest that the *shadow-1* and *shadow-2* mutants have enhanced shade tolerance and a better ability to maintain tiller numbers under low-light conditions than wild-type plants.

**Figure 2:**
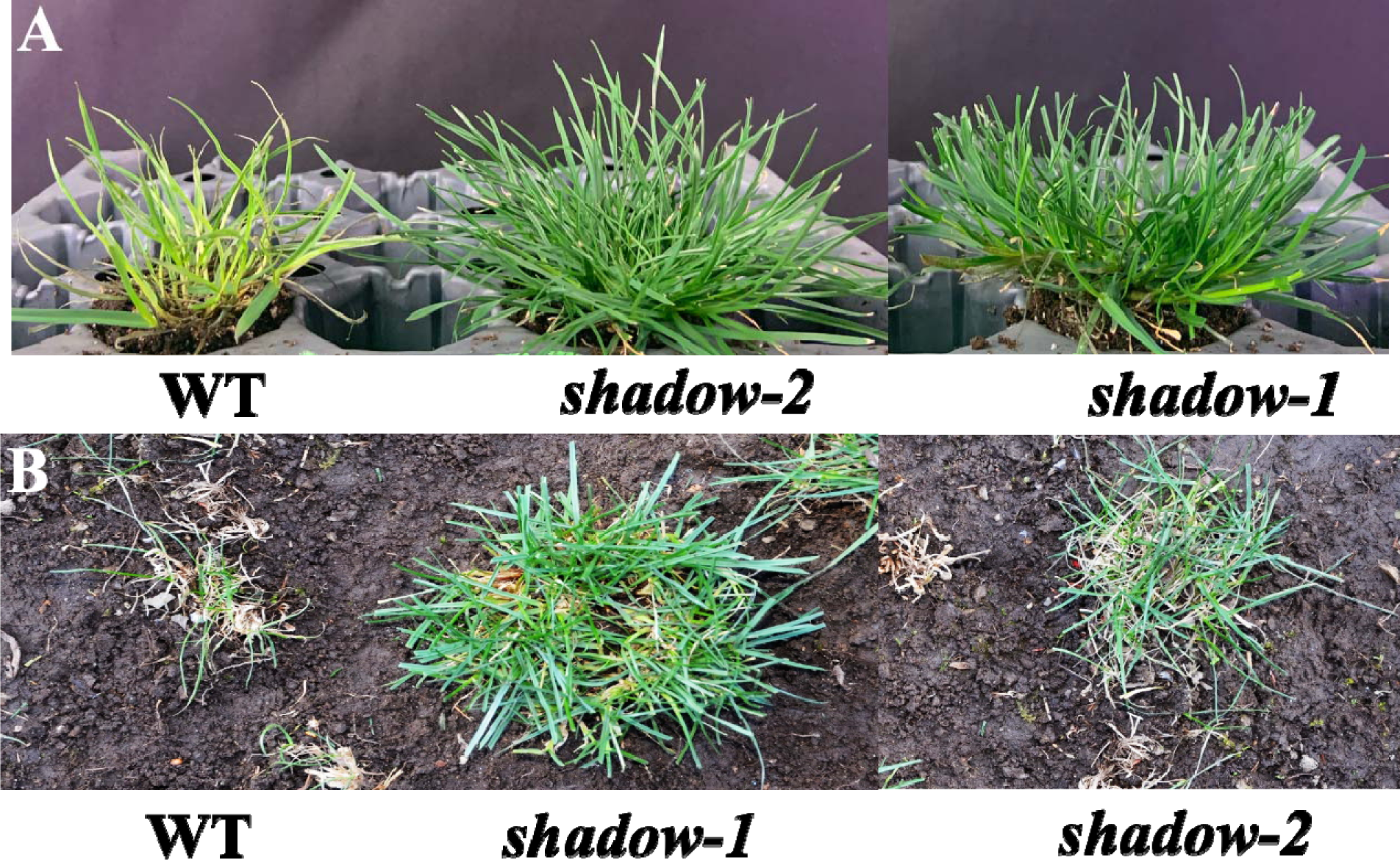
Growth and health of *shadow-1* and *shadow-2 m*utants under shade conditions. **(A).** Growth of *shadow-1* and *shadow-2* mutants under 95% controlled shade conditions in a greenhouse for four weeks. The wild-type control plants show reduced growth, fewer tillers, and yellowing of leaves, indicating suboptimal growth under the shade conditions (left). The *shadow-2* mutant exhibits a healthy canopy with dense and green foliage under the same shade conditions (middle). The *shadow-1* mutant shows a healthy canopy with dense and green foliage, performing better overall compared to *shadow-2* under the same shade conditions (right). (**B).** Field performance of *shadow-1* and *shadow-2* mutants under approximately 85% natural shade conditions in the woods for 30 months. The wild-type control plants are diminishing (right). The *shadow-1* mutant plants have healthy green foliage, indicating good adaptation to 85% natural shade conditions (middle). The *shadow-2* mutant plants show intermediate growth performance with some yellowing and reduced tillers (right).

A greenhouse experiment was conducted to investigate the effect of shade on tillering in wild-type, *shadow-1,* and *shadow-2* plants. After six weeks of growth under light conditions, plants were transferred to 95% shade. At the start of the shade treatment, *wild-type*, *shadow-1,* and *shadow-2* plants had an average of 15.2, 22.4, and 21.4 tillers per plug, respectively. After four weeks, light-grown plants had a significant increase in tiller numbers of 6.9, 14.3, and 10.8 in wild-type, *shadow-1*, and *shadow-2,* respectively. However, shade-treated plants had a non-significant increase in tiller number, with only 0.7, 1.1, and 0.6 tillers per plug in wild-type, *shadow-1*, and *shadow-2*, respectively (Figure 3, 2A). These results demonstrate that shade inhibits tillering in both wild-type and shade-tolerant mutants.

**Figure 3.**
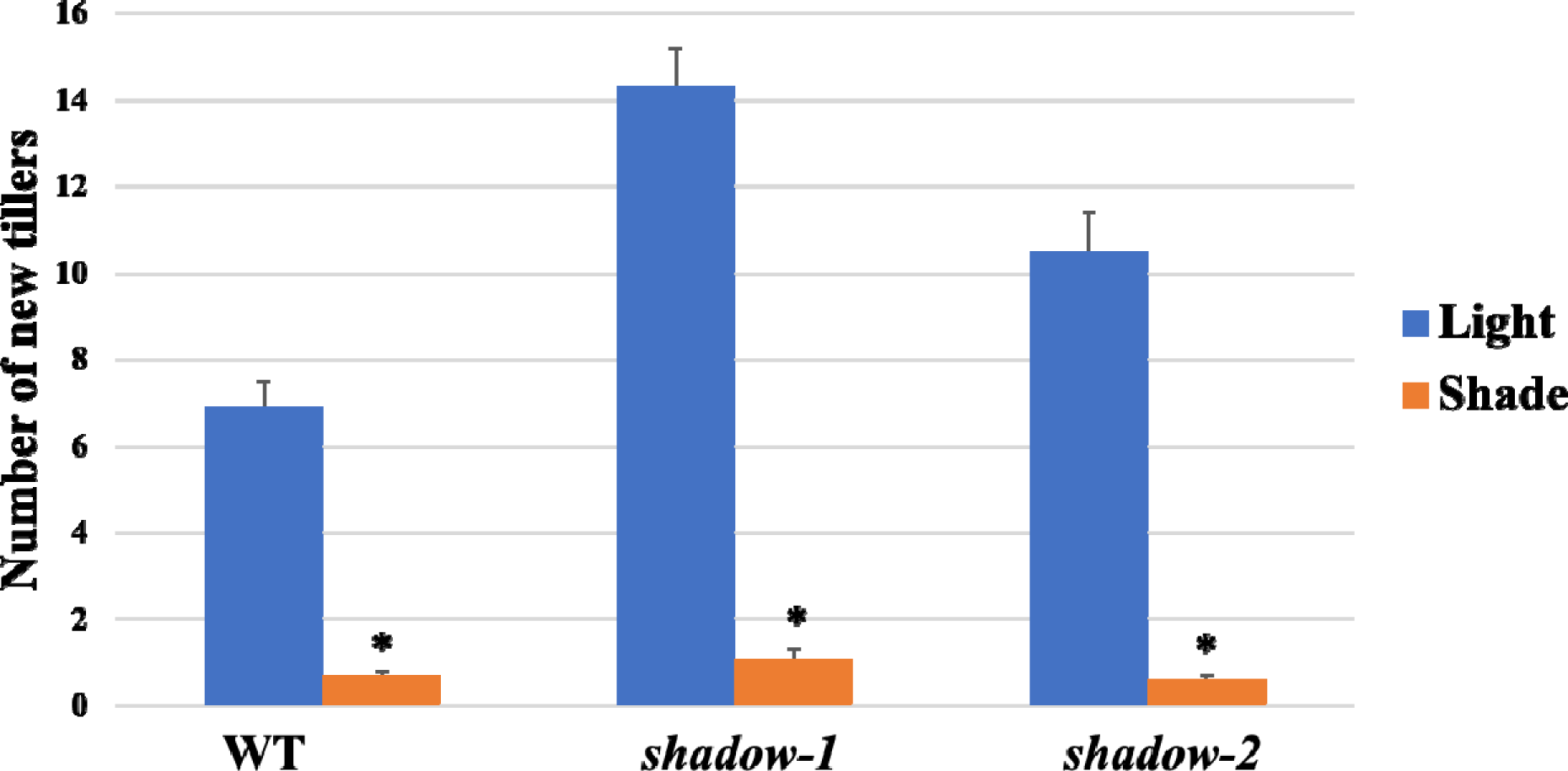
Deep shade-mediated inhibition of tillering in wild-type, *shadow-1*, and *shadow-2* plants. Plants with five tillers were grown under full light for six weeks. Subsequently, they were transferred to both full light and 95% shade for a period of four weeks, and new tiller numbers were recorded after the end of the treatment. Under full light, new tillers were observed in wild-type and both mutants, while under the shade conditions, there were no significant new tiller observed in wild-type and both mutants. * Mean differing from light conditions at P < 0.001.

### 3.3. Possible role of CKs and ABA hormones in high tillering in *shadow-1* and *shadow-2* mutants

In this study, we investigated the role of plant hormones in regulating tillering by measuring hormone levels in wild-type and mutant plants. Our results revealed significant differences in the levels of CKs and ABA between wild-type and mutants. CKs, known to be positive regulators of tillering, were found to be 2.9- and 1.9-fold higher in *shadow-1* and *shadow-2* mutants, respectively, compared to the wild-type (Figure 4B). Conversely, the levels of ABA, a negative regulator of tillering, were found to be 1.5-fold lower in both mutants compared to the wild-type (Figure 4D). Our findings suggest that the altered levels of CKs and ABA in the mutants might play important roles in more tillering.

**Figure 4.**
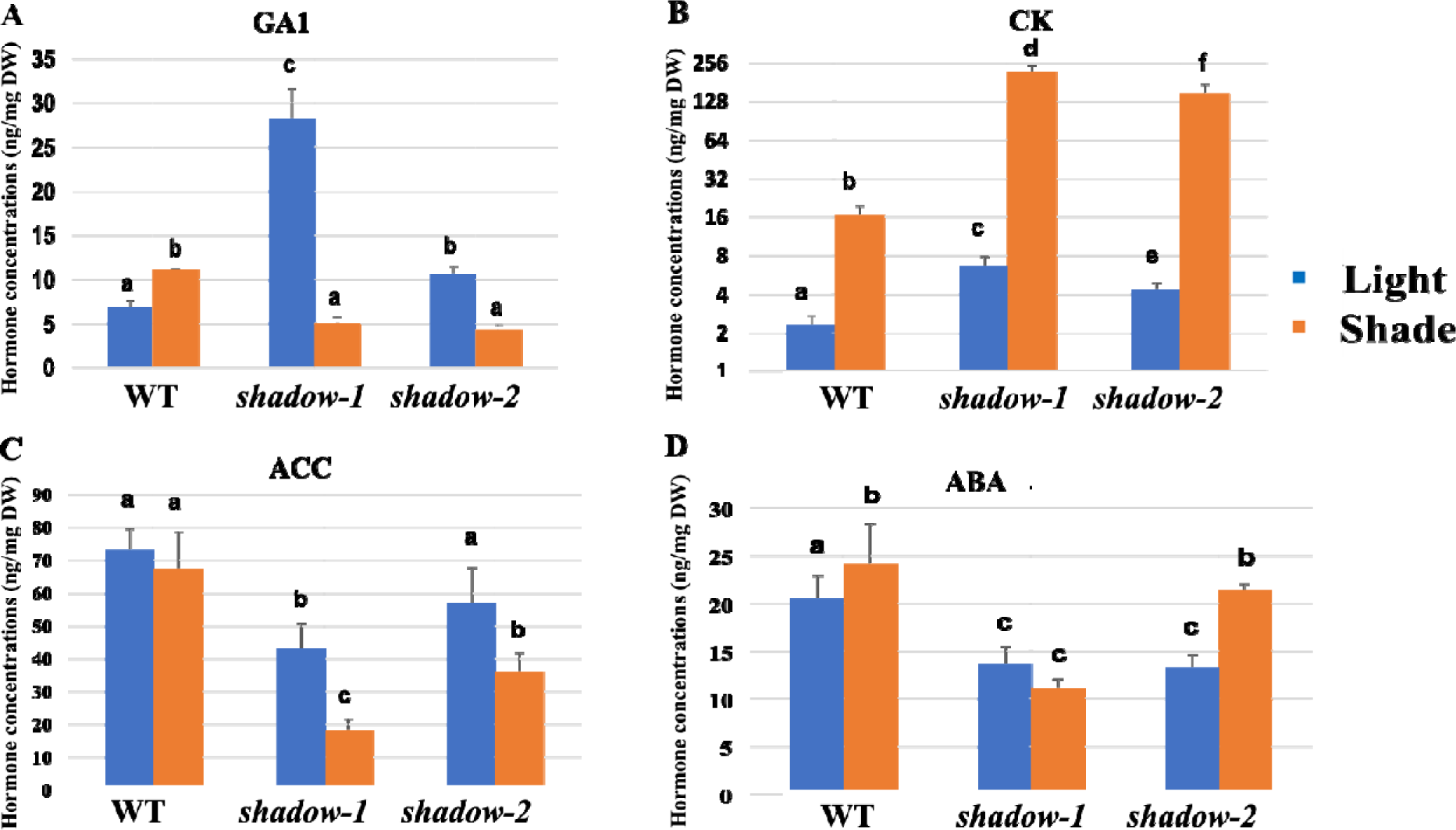
Plant hormone concentrations in wild-type, *shadow-1*, and s*hadow-2* mutant plants under light and shade conditions. **(A).** GA1 content was higher in both mutants compared to wild-type under light conditions. Under 95% shade conditions, GA1 content in the wild-type plants was elevated, but in both mutants, their GA1 levels were reduced when compared to the same types of plants under light conditions. **(B).** Significantly higher concentrations of CKs were observed in both mutants compared to wild-type under both light and shade conditions. **(C).** Reductions in ACC (a key ethylene precursor) were observed in both mutants compared to the wild-type plants under light and shade conditions. **(D).** Under the light conditions, both mutants had lower concentrations of ABA compared to wild-type. While under shade conditions, ABA concentrations were increased in both wild-type and *shadow-2 mutant*. *Shadow-1* mutant has lower ABA levels under both light and shade conditions. Data represent the average of three replicates for each sample. Each replicate consists of the pooled leaf samples from 10 plants. Bars indicate standard deviations. Values followed by the same letter were not significantly different from each other, according to the least significant difference (LSD) (P < 0.05).

**Figure 5.**
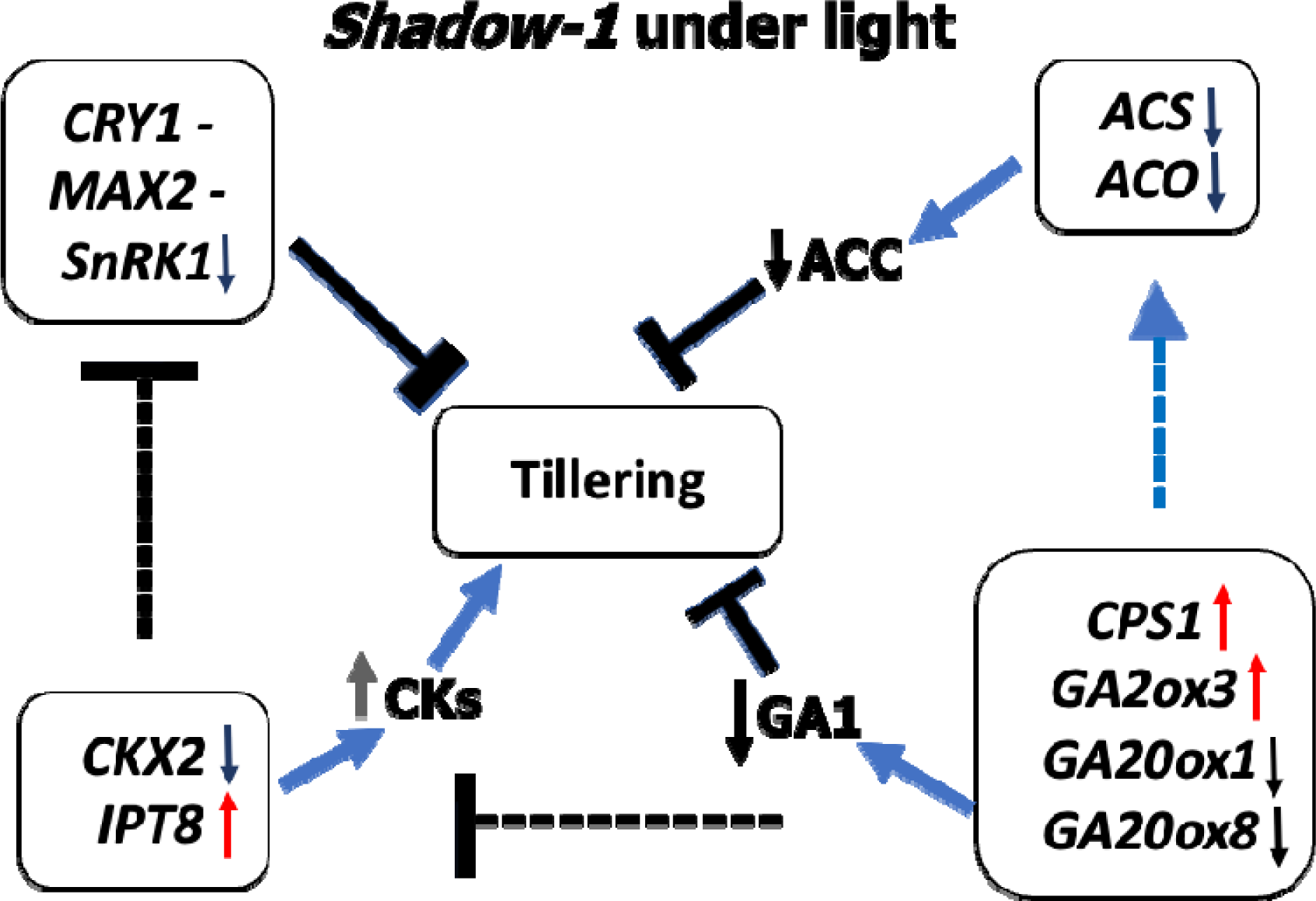
A summary of changes in expression of tillering-related genes observed in the *shadow-1* mutant of perennial ryegrass compared to the wild-type counterpart under light conditions. Down-regulation of *SnRK1* and *CKX2*, and up-regulation of *IPT8* gene *shadow-1* mutant compared to wild-type in might be involved in high tillering. ***CRY1* (**cryptochrome 1): Tillering inhibitor; ***MAX2*** (more axillary growth 2): Tillering inhibitor; ***SnRK1*** (sucrose non-fermenting kinase): Tillering inhibitor; ***CKX2*** (cytokinin oxidase/dehydrogenase 2): Cytokinin degradation; ***IPT8*** (isopentenyl transferase 8): Cytokinin biosynthesis; ***ACS*** (aminocyclopropanecarboxylate synthese): Ethylene biosynthesis; ***ACO*** (aminocyclopropanecarboxylate oxidase): Ethylene biosynthesis; ***CPS1*** (ent-copalyl diphosphate synthase): Gibberellic acid biosynthesis; ***GA2ox3*** (GA2 oxidase 3): Gibberellic acid biosynthesis; ***GA20ox1*** (GA20 oxidase 1): Gibberellic acid biosynthesis; ***GA20ox8*** (GA20 oxidase 8): Gibberellic acid biosynthesis; Arrow, T-bars, and hyphen indicate upregulation, downregulation, and no changes, respectively.

To further understand the molecular mechanisms underlying the regulation of tillering in the *shadow-1* mutant, we conducted transcriptome sequencing and compared the gene expression profiles between wild-type and *shadow-1* under light conditions. The results showed that the expression levels of genes related to CKs, and ABA hormone regulation were differentially expressed between the two groups, consistent with the hormonal data (Table 2). In particular, the expression level of the CKs negative regulator *CKX2* gene was significantly decreased in the *shadow-1* mutant compared to the wild-type (15.4-fold), while the expression level of *IPT8* gene, which promotes CK synthesis, was 4.6-fold higher in the *shadow-1* mutant. Moreover, the *GA2ox8* gene, which regulates ABA level, was upregulated 2.8-fold in the *shadow-1* mutant compared to the wild-type. In addition, *SnRK1*, a negative regulator of tillering in plants, was found to be significantly decreased in expression level by 3.8-fold in the *shadow-1* mutant compared to wild-type. These findings provide further evidence that CKs and ABA hormones together might play important roles in the high-tillering phenotypes observed in the *shadow-1* mutant.

**Table 2:**
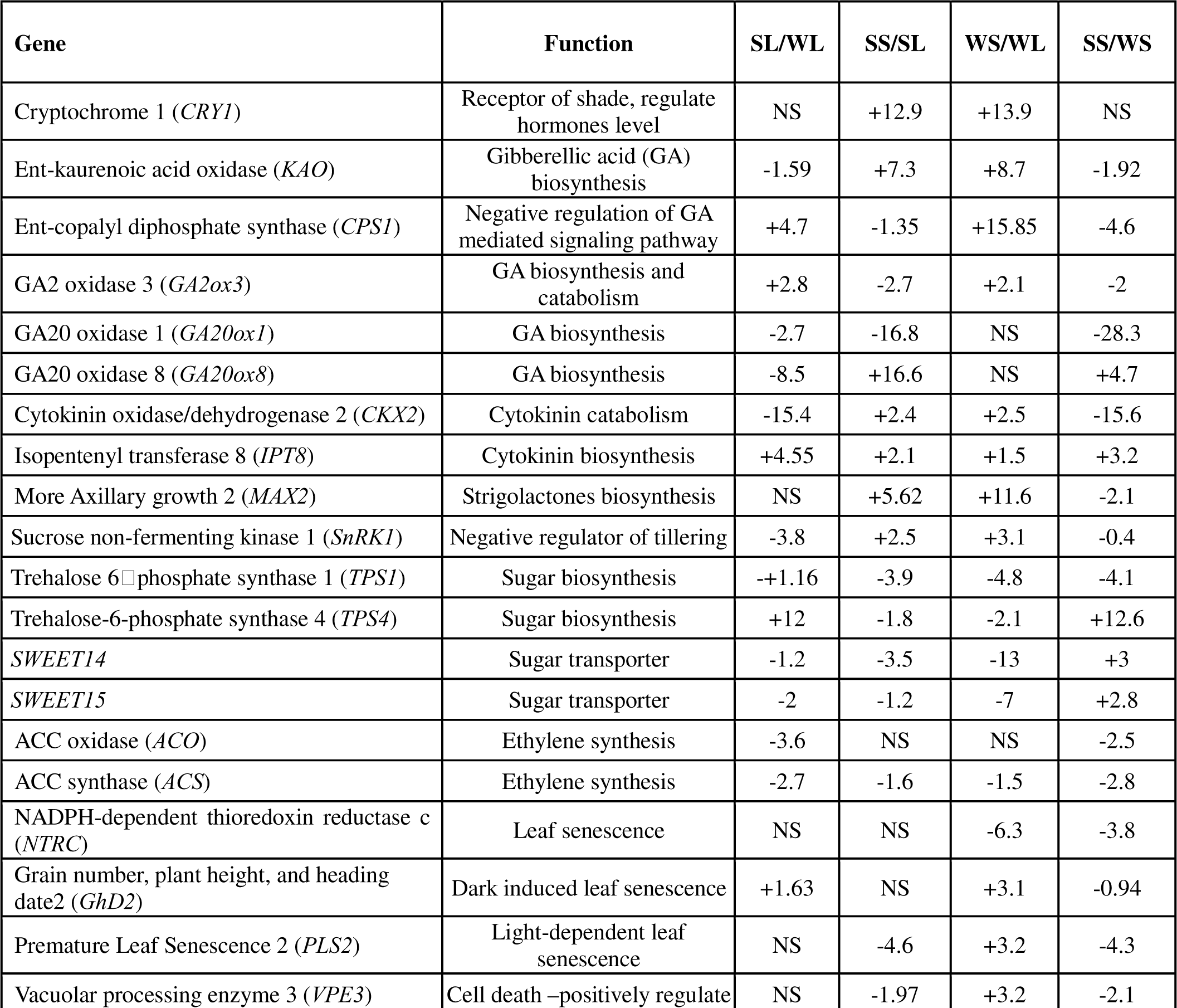
A list of differentially expressed genes between *shadow-1* mutant and wild-type under light and shade conditions. Differential gene expression was calculated based on the fold change (relative expression) compared to the wild-type control. “-” represent down-regulation of the genes and “+” represent up-regulation of the genes. NS denotes no significant differences between the samples. SL and WL denotes *shadow-1* mutant and wild-type under light conditions, respectively. Whereas, SS and WS denotes *shadow-1* mutant and wild-type under shade conditions, respectively.

| Gene | Function | SL/WL | SS/SL | WS/WL | SS/WS |
| --- | --- | --- | --- | --- | --- |
| Cryptochrome 1 (*CRY1*) | Receptor of shade, regulate hormones level | NS | +12.9 | +13.9 | NS |
| Ent-kaurenoic acid oxidase (*KAO*) | Gibberellic acid (GA) biosynthesis | -1.59 | +7.3 | +8.7 | -1.92 |
| Ent-copalyl diphosphate synthase (*CPS1*) | Negative regulation of GA mediated signaling pathway | +4.7 | -1.35 | +15.85 | -4.6 |
| GA2 oxidase 3 (*GA2ox3*) | GA biosynthesis and catabolism | +2.8 | -2.7 | +2.1 | -2 |
| GA20 oxidase 1 (*GA20ox1*) | GA biosynthesis | -2.7 | -16.8 | NS | -28.3 |
| GA20 oxidase 8 (*GA20ox8*) | GA biosynthesis | -8.5 | +16.6 | NS | +4.7 |
| Cytokinin oxidase/dehydrogenase 2 (*CKX2*) | Cytokinin catabolism | -15.4 | +2.4 | +2.5 | -15.6 |
| Isopentenyl transferase 8 (*IPT8*) | Cytokinin biosynthesis | +4.55 | +2.1 | +1.5 | +3.2 |
| More Axillary growth 2 (*MAX2*) | Strigolactones biosynthesis | NS | +5.62 | +11.6 | -2.1 |
| Sucrose non-fermenting kinase 1 (*SnRK1*) | Negative regulator of tillering | -3.8 | +2.5 | +3.1 | -0.4 |
| Trehalose 6-phosphate synthase 1 (*TPS1*) | Sugar biosynthesis | -+1.16 | -3.9 | -4.8 | -4.1 |
| Trehalose-6-phosphate synthase 4 (*TPS4*) | Sugar biosynthesis | +12 | -1.8 | -2.1 | +12.6 |
| *SWEET14* | Sugar transporter | -1.2 | -3.5 | -13 | +3 |
| *SWEET15* | Sugar transporter | -2 | -1.2 | -7 | +2.8 |
| ACC oxidase (*ACO*) | Ethylene synthesis | -3.6 | NS | NS | -2.5 |
| ACC synthase (*ACS*) | Ethylene synthesis | -2.7 | -1.6 | -1.5 | -2.8 |
| NADPH-dependent thioredoxin reductase c (*NTRC*) | Leaf senescence | NS | NS | -6.3 | -3.8 |
| Grain number, plant height, and heading date2 (*Ghd2*) | Dark induced leaf senescence | +1.63 | NS | +3.1 | -0.94 |
| Premature Leaf Senescence 2 (*PLS2*) | Light-dependent leaf senescence | NS | -4.6 | +3.2 | -4.3 |
| Vacuolar processing enzyme 3 (*VPE3*) | Cell death –positively regulate | NS | -1.97 | +3.2 | -2.1 |

### 3.4. Tillering-inhibitor genes overexpressed under shade conditions

Previous studies have identified *MAX2*, *CRY1*, and *SnRK1* genes as negative regulators of tillering in plants (Stirnberg *et al*. 2007; Minakuchi *et al*. 2010; Tsai and Gazzarrini, 2014; Zhai *et al*. 2020). To investigate the potential role of these genes in the shade-induced inhibition of tillering, we measured their expression levels in wild-type and *shadow-1* plants under shade and light conditions. The results showed that shade treatment led to significant increases in the expression levels of *MAX2*, *SnRK1*, and *CRY1* genes in both wild-type and *shadow-1* plants (Table 2). *MAX2* gene expression was elevated by 11.6- and 5.6-fold in wild-type and *shadow-1* plants, respectively, under shade conditions compared to light conditions. *SnRK1* gene expression was up-regulated by 2.5- and 3.1-fold in wild-type and *shadow-1* plants, respectively, under shade conditions compared to light conditions. Similarly, *CRY1* gene expression was up- regulated by 13.9- and 12.9-fold in wild-type and *shadow-1* plants, respectively, under shade conditions compared to wild-type plants (Table 2). These results suggest that the shade-induced upregulation of *MAX2*, *CRY1*, and *SnRK1* gene expression might be involved in the inhibition of tillering under shade conditions (Figure 7).

### 3.5. Possible role of GA, CKs, ETs, and ABA hormones in shade tolerance in *shadow-1* and shadow-2 mutants

The study aimed to investigate the differential responses of wild-type and mutants to shade stresses by examining hormonal profiles (Figure 4). One of the major symptoms that plants exhibit during shade avoidance response is shoot elongation, which requires GA under shade conditions (Pierik *et al*. 2004a). Our results revealed that the bioactive GA1 level increased 1.6-fold in wild-type under shade conditions compared to light conditions (Figure 4A). In contrast, GA level decreased 5.6- and 2.5-fold in *shadow-1* and *shadow-2* mutants, respectively, under shade conditions compared to light. Furthermore, both mutants exhibited a significantly reduced shade avoidance response (shoot elongation) compared to wild-type under shade conditions on the field, with a reduction of 2.2- and 2.6-fold in *shadow-1* and *shadow-2*, respectively. These findings suggest that GA plays a crucial role in shade tolerance, as a reduced shade avoidance response was observed in both mutants due to the absence of GA1.

In addition to GA, the results showed the potential role of ET in shade tolerance in both mutants. It was found that the ACC level was 2.4- and 1.6-fold lower in *shadow-1* and *shadow-2* mutants, respectively, under shade conditions compared to light, while there was no significant change in ACC level in the wild-type. Furthermore, *shadow-1* and *shadow-2* mutants exhibited reductions of 3.7- and 1.9-fold, respectively, in ACC level compared to the wild-type under shade conditions (Figure 4C). Additionally, we observed that ABA levels were reduced by 2.2- and 1.13-fold in *shadow-1* and *shadow-2* mutants, respectively, compared to the wild-type under shade conditions (Figure 4D).

Previous studies have shown that CKs are positive regulators of shade tolerance in plants, as they enhance shade tolerance and delay senescence under shade conditions (Xing *et al*. 2009). In this study, we found that the CKs level was increased by 7.3-, 32.6-, and 33.6-fold in the wild-type, *shadow-1,* and *shadow-2,* respectively, under shade conditions compared to light conditions. Moreover, we observed that the CKs level was increased by 13.1- and 8.9-fold in *shadow-1* and *shadow-2* mutants, respectively, compared to the wild-type under shade conditions (Figure 4B). These results suggest that CKs may be involved in the shade tolerance of *shadow-1* and *shadow-2* mutants. These findings suggest that reductions in GA and ET synthesis, along with an increase in CKs synthesis, may contribute to the shade tolerance observed in *shadow-1* and *shadow-2* mutants.

### 3.6. Transcriptome data suggest the role of GA, sugar, and senescence regulating genes in shade tolerance in *shadow-1*

The response of plants to shade is characterized by various changes in their morphology and physiology, including tillering inhibition, chlorophyll degradation, shoot elongation, and senescence. To investigate the molecular mechanisms underlying these responses, we analyzed the differential expression of genes involved in these processes under shade conditions compared to light (Table 2). Our results showed that GA biosynthesis and metabolism genes are differentially expressed between wild-type and *shadow-1* mutant plants in response to shade. *KAO, CPS1,* and *GA2ox3* genes were up-regulated 8.7-, 15.9-, and 2.1-fold, respectively, in wild-type plants under shade conditions compared to light conditions (Table 2, Figure 7A). Conversely, GA biosynthesis genes, *KAO, CPS1*, and *GA20ox1*, were down-regulated by 1.35-, 2.7-, and 16.8-fold, respectively, in *shadow-1* mutant plants under shade conditions compared to light conditions (Table 2, Figure 7B). Furthermore, *GA20ox8*, a negative regulator of GA, was up-regulated by 16.6-fold in the *shadow-1* mutant under shade conditions compared to light (Table 2, Figure 7B). In contrast, there was no significant change in *GA20ox8* expression levels in the wild-type under shade conditions. The findings elucidated the molecular mechanism behind the decreased GA1 levels observed in the *shadow-1* mutant in comparison to the wild-type when subjected to shade conditions. Furthermore, they provided evidence for the involvement of GA in enhancing shade tolerance in the *shadow-1* mutant.

Senescence-related genes are differentially expressed in response to shade in both the wild-type and *shadow-1* mutant (Table 2). Under shade conditions, the Grain number, plant height, and heading date2 (*GhD2*) gene was up-regulated by 3.1-fold in the wild-type compared to light conditions, while no change was observed in the *shadow-1* mutant. Conversely, NADPH-thioredoxin reductase, type C (*NTRC*) gene was down-regulated by 6.1-fold in the wild-type compared to light conditions, but no change was observed in the *shadow-1 mutant*. Additionally, premature leaf senescence (*PLS2*) gene was up-regulated by 3.2-fold in the wild-type under shade conditions compared to light, while it was down-regulated by 4.6-fold in the *shadow-1* mutant. Similarly, vacuolar processing enzyme (*VPE3*) gene was up-regulated by 3.2-fold in the wild-type under shade conditions compared to light conditions, while it was down-regulated by 2-fold in the *shadow-1* mutant. These findings suggest that the differential expression of these senescence-related genes may contribute to the observed differences in shade response between the wild-type and *shadow-1* mutant.

Ribulose-1,5-bisphosphate carboxylase-oxygenase (Rubisco) gene is one of the most important gene of photosynthesis. Rubisco gene was 3.5 and 1.3-fold down-regulated in wild-type and *shadow-1*, respectively, under shade conditions compared to light conditions (Table 2). Sugar biosynthesis genes were differentially expressed between the wild-type and *shadow-1* mutant under light and shade conditions. Sugar biosynthesis genes, trehalose 6 phosphate synthase 1 (*TPS1*) and trehalose-6-phosphate synthase 4 (*TPS4*), were down-regulated in both wild-type and *shadow-1* under shade conditions compared to light conditions. However, the reduction of this expression in the *shadow-1* mutant was lower than the one in wild-type (Table 2). The expression level of *TPS4* gene was 12.6-fold higher in *shadow-1* mutant compared to wild-type under shade conditions. Two sugar transporter genes, *SWEET14* and *SWEET15*, were 13- and 7-fold, respectively, downregulated in wild-type under shade conditions compared to light conditions, while these genes 3.5- and 1.2-fold, respectively, downregulated in *shadow-1* mutant under shade conditions compared to light conditions.

### 3.7. Validation of RNA-Seq data by qRT**□**PCR

To validate the transcriptome data, four important differentially expressed genes were selected for gene expression analysis via qRT PCR (Figure 6A). Similar to transcriptome data, *CKX2* and *SnRk1* genes were downregulated in the *shadow-1* compared to wild-type under light conditions. Congruent with transcriptome data, a significantly overexpression of *MAX2* and *SnRk1* genes were observed in the wild-type under the shade conditions compared to light conditions (Figure 6B). Overall, the qRT-PCR results validated the expression pattern of differentially expressed genes observed in transcriptome data.

**Figure 6.**
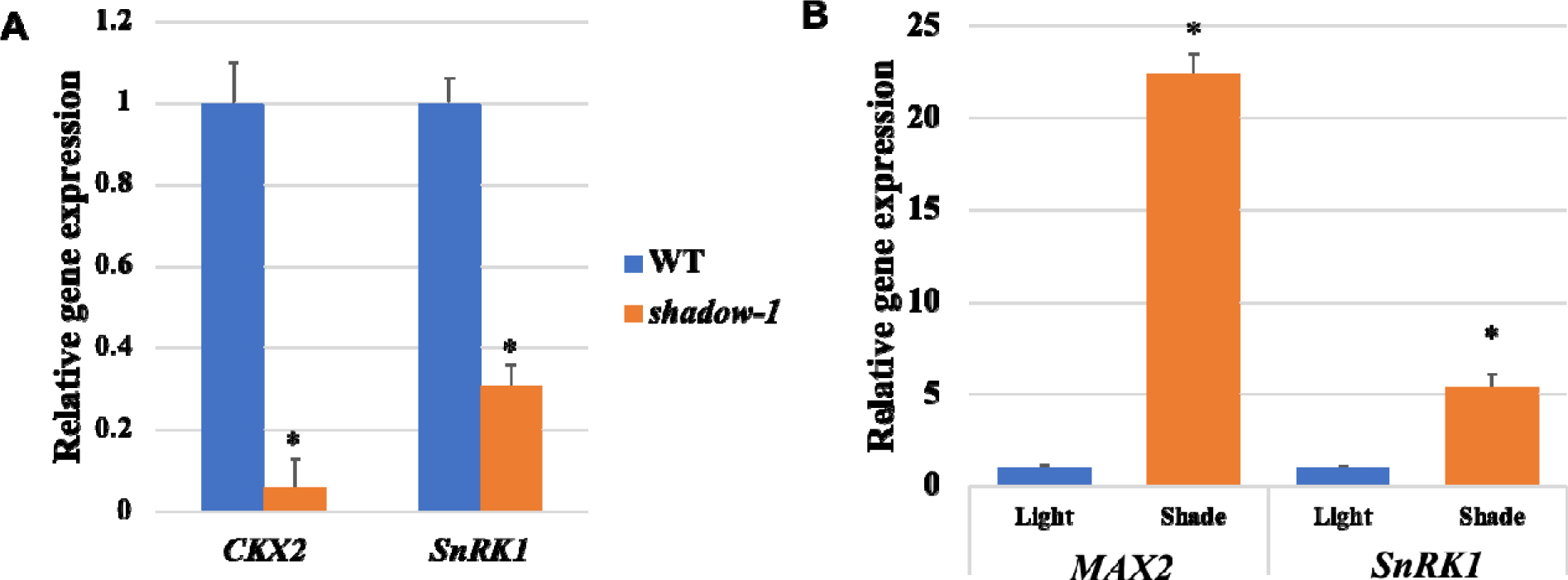
qRT-PCR data verified the accuracy of transcriptome analysis. **(A).** Relatively lower expression levels of *CKX2* and *SnRK1* genes in *shadow-1* compared to wild-type plants were observed by qRT-PCR. **(B).** Relatively higher expression levels of *MAX2* and *SnRK1* gene in wild-type under light compared to shade conditions. Gene expression levels in each sample were normalized using the expression level of the internal control, *LpGAPDH*, in the sam sample. The changes in the gene expression level were similar in qRT-PCR and RNAseq data. The data presented are the averages of three independent technical replicates. Bars indicate standard deviations. Asterisks (*) represent a significant difference in the gene expression levels in *shadow-1* mutant in comparison to compared to wild-type, using a two-tailed Student’s t-test (*P* < 0.05).

**Figure 7.**
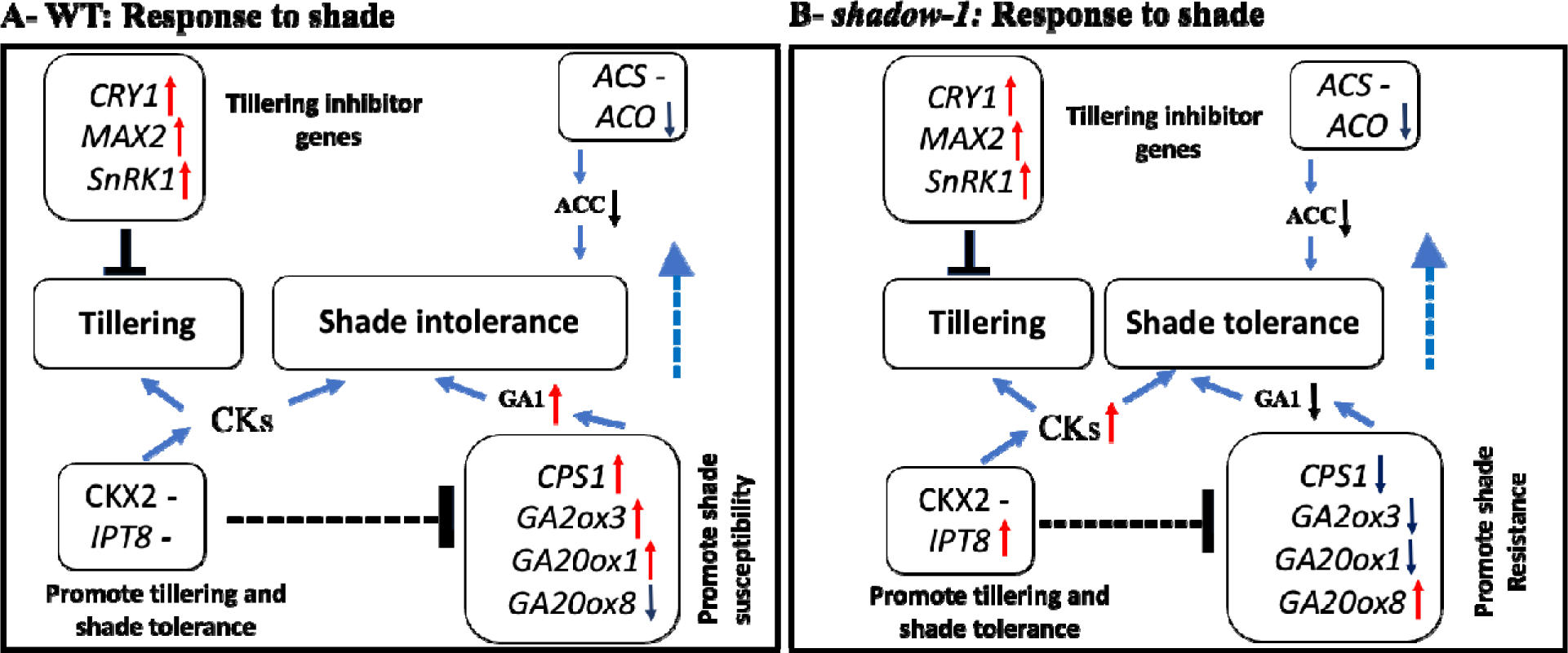
A summary of changes in expression of tillering and shade tolerance-related gene observed in WT and *shadow-1* mutant of perennial ryegrass under deep (95%) shade conditions. **(A) WT:** In wild-type, tillering inhibitor genes, *CRY1*, *MAX2,* and *SnRK1* and GA biosynthesis genes *CPS1*, *GA2ox3*, and *GA20ox1* genes were upregulated under shade conditions compared to light conditions, which leads to tillering inhibition and shade susceptibility. **(B**) ***shadow-1*; Shade Vs. Light Conditions** In *shadow-1,* tillering inhibitor genes and cytokinin biosynthesis gene *IPT8* were upregulated in *shadow-1* mutant under shade conditions compared to light conditions, conversely, GA biosynthesis genes, *CPS1*, *GA2ox3*, and *GA20ox1* were down-regulated. Altogether these changes lead to tillering inhibition and shade tolerance in *shadow-1* mutant. ***CRY1* (**cryptochrome 1): Tillering inhibitor; ***MAX2*** (more axillary growth 2): Tillering inhibitor; ***SnRK1*** (sucrose non-fermenting kinase): Tillering inhibitor; ***CKX2*** (cytokinin oxidase/dehydrogenase 2): Cytokinin degradation; ***IPT8*** (isopentenyl transferase 8): Cytokinin biosynthesis; ***ACS*** (aminocyclopropanecarboxylate synthese): Ethylene biosynthesis; ***ACO*** (aminocyclopropanecarboxylate oxidase): Ethylene biosynthesis; ***CPS1*** (ent-copalyl diphosphate synthase): Gibberellic acid biosynthesis; ***GA2ox3*** (GA2 oxidase 3): Gibberellic acid biosynthesis; ***GA20ox1*** (GA20 oxidase 1): Gibberellic acid biosynthesis; ***GA20ox8*** (GA20 oxidase 8). Arrow, T-bars, and hyphen indicate promotion, inhibition, and no changes, respectively.

## 4. Discussion

In this study, we present findings on changes in tillering, shade tolerance, hormonal levels, and gene expression in wild-type and dwarf mutant plants of perennial ryegrass. Our results show that both *shadow-1* and *shadow-2* mutants exhibit significantly higher levels of tillering in light conditions compared to their wild-type counterparts. Additionally, both mutants demonstrated shade tolerance in plant height and canopy color under shade conditions. However, like the wild-type plants, both mutants also display shade-induced inhibition of tillering under 95% shade conditions. We further demonstrate that under 95% shade conditions, wild-type plants exhibit increased GA and reduced CK levels, while *shadow-1* mutant plants have reduced GA but increased CK levels. This explains why the *shadow-1* mutant is shade tolerant in terms of plant height, grass quality, and color. Conversely, we observed that the tillering inhibition-related genes *CRY1*, *MAX2*, and *SnRK1* are upregulated in both wild-type and *shadow-1* mutant plants, providing molecular evidence for the inhibition of tillering under 95% shade conditions in the *shadow-1* mutant. These results suggest that the shade tolerance observed in plant height and leaf color in *shadow-1* is not linked with tillering under deep shade conditions, contributing to our understanding of the mechanisms behind tillering and shade tolerance in turfgrasses.

Various modern and tradition methods are used to create useful variations in plants for molecular study and plant breeding (Kumar *et al*. 2013, Rana *et al*. 2022, Kumar *et al*. 2023a, Liu *et al*. 2023, Kumar *et al*. 2023b, Kumar and Yer 2023, Chugh *et al*. 2024, Kumar *et al*. 2024). Mutagenesis is one of the most powerful methods for creating disable variations (Kumar et al. 2022, Li et al, 2016, Li et al, 2017). The critical role of CKs in influencing tiller numbers is well-established, with increased levels of endogenous CKs known to promote tiller formation (Zalewski *et al*. 2010; Yeh *et al*. 2015; Wang *et al*. 2018; Liu *et al*. 2019). The *CKX2* gene, reported to down-regulate CKs, when silenced, results in enhanced endogenous CKs and promotes tiller formation in rice and barley (Zalewski *et al*. 2010; Yeh *et al*. 2015; Zhuang *et al*. 2019). The decreased expression of the *CKX2* gene coupled with increased cytokinin levels in our mutant plants illustrate their contribution to the augmentation of tillering under light-grown conditions. Prior studies have also reported the negative role of ABA hormone in tillering (Wang *et al*. 2018; Liu *et al*. 2019). In our current research, we provide evidence that the *shadow-1* and *shadow-2* mutants contain lower levels of ABA compared to the wild-type. These findings propose that ABA may be involved in the tillering process of perennial ryegrass.

*SnRK1* is a crucial regulator of energy homeostasis in plants, closely associated with plant growth and development (Tsai and Gazzarrini, 2014). Previous research has demonstrated the negative regulation of tillering by *SnRK1* in plants (Tsai and Gazzarrini *et al*. 2014; Simon *et al*. 2018; Baena *et al*. 2020; Belda *et al*. 2020). *SnRK1* is involve activated by various stresses in plants (Mao *et al*. 2020). *SnRK* gene mediate crosstalk between ABA and CKs signaling. GA positively regulates the expression of *SnRK1* gene in plants (Li *et al*. 2023). Application of GA biosynthesis inhibitor, Paclobutrazol, increases CKs and suppresses the of *SnRK1* gene in plants (Li *et al*. 2023). In our study, we observed lower expression of the *SnRK1* gene in the *shadow-1* mutant compared to wild-type. Filipe *et al*. (2018) have reported that overexpression of *OsSnRK1* gene in rice reduce 40% tiller compared to wild-type. Similarly, Zhang *et al*. (2023) demonstrated that the overexpression of the wheat *TaSnRK1* gene in rice leads to a decrease in tiller number compared to the wild-type under field conditions. Rice ubiquitin conjugating enzyme *OsUbc13* negatively regulates *OsSnRK1* gene and downregulation of *OsUbc13* gene via RNAi decrease tiller number in rice (Liu *et al*. 2023). An increase in CKs hormone in both mutants and downregulation of *SnRK1* gene in the *shadow-1* mutant suggest their involvement in high tillering phenotype.

We found that shade inhibits tillering in both wild-type and mutants, though both mutants showed more tolerance to the shade stress than the wild-type plants. The results reveal that the negative tillering regulators *MAX2, CRY1,* and *SnRK1* genes were up-regulated under shade conditions. The *MAX2* gene has been shown to be up-regulated under low light conditions in several plant species, such as sorghum and *Arabidopsis* (Shen *et al*. 2007; Kebrom *et al*. 2016). In sorghum, shade treatment up-regulates the *MAX2* gene, which inhibits bud outgrowth. Congruent with these studies, our study showed an elevated expression of the *MAX2* gene under shade conditions in perennial ryegrass, indicating a negative role of this gene in tillering. Cryptochrome is a photoreceptor that plays a crucial role in receiving shade signals in plants (Zhai *et al*. 2020; Lyu *et al*. 2021). Cryptochrome regulates shade avoidance responses, including shoot elongation through increasing GA level in plants. In *Arabidopsis*, Cryptochrome 1 has been found to inhibit shoot branching, suggesting a negative role in tillering. In our study, the *CRY1* gene was highly elevated under shade conditions in both wild-type and *shadow-1* mutant plants, suggesting a possible role of *CRY1* in shade-induced tillering inhibition in perennial ryegrass. Similarly, the *SnRK1* gene, a negative regulator of tillering, was up-regulated under shade conditions in our study, suggesting the role of *SnRK1* in shade-induced tillering inhibition. Our findings highlight the possible role of negative regulators of tillering, *MAX2, CRY1*, and *SnRK1* genes, in the shade-induced inhibition of tillering.

The hormone data obtained in this study suggests that GA, CKs, and ET may be involved in the shade tolerance mechanisms of the *shadow-1* and *shadow-2* mutants. These hormones have been reported to play a role in shade tolerance in previous studies (Pierik *et al*. 2004a; Pierik *et al*. 2004b; Das *et al*. 2016; Li *et al*. 2017, Zhang *et al*. 2021). Elevated concentrations of gibberellins have been observed in shade-treated beans (Beall *et al*. 1996), sunflowers (Kurepin *et al*. 2007), and *Arabidopsis* (Bou-Torrent *et al*. 2014; Liu *et al*. 2016). In plants, the expression of GA biosynthesis genes is induced by shade treatment, which increases the level of GA (Hisamatsu *et al*. 2005; Yu *et al*. 2015; Li *et al*. 2017). Pierik *et al*. (2004) reported that inhibiting gibberellin production in tobacco increases shade tolerance by restricting shoot elongation. In a transcriptome analysis of the *shadow-1* mutant, Li *et al*. (2017) found that the GA biosynthesis pathway was downregulated compared to the wild-type under shade conditions, suggesting that GA hormones may play a role in shade tolerance. Shade treatment induces ET production in wild-type tobacco, which results in a shade avoidance syndrome. In contrast, ET-insensitive tobacco exhibits shade tolerance with reduced levels of ET (Pierik *et al*. 2004a). In this study, we observed a reduction in ACC (ET) levels in both mutants under shade conditions compared to light, while the ACC level did not change in the wild-type, indicating a possible role of ET in shade tolerance in mutants. Elevated production of CKs in plants has been reported to increase shade tolerance and delay senescence (Gan *et al*. 1995; Robson *et al*. 2004; Xing *et al*. 2009; Xu *et al*. 2010; Schafer *et al*. 2015). Overexpression of the *IPT* transgene in creeping bentgrass and perennial ryegrass increases endogenous CKs content, resulting in increased shade tolerance and delayed senescence (Robson *et al*. 2004; Xu *et al*. 2010). In this study, we observed an elevated level of CKs in both mutants compared to the wild-type under light and shade conditions, suggesting a role for CKs in shade tolerance. Taken together, the results of our study suggest that CKs, GA, and ET hormones may play roles in the shade tolerance mechanisms of both mutants. These findings are consistent with previous studies and provide further insights into the complex hormonal regulation of shade tolerance in plants.

The transcriptome results of our study suggest that the shade tolerance of the *shadow-1* mutant is associated with the differential expression of senescence-related genes compared to wild-type under shade treatment. The *GhD2, NTRC, PLS2*, and *VPE3* genes showed different expression patterns in the *shadow-1* mutant compared to the wild-type under shade treatment. The *GhD2* gene was up-regulated in the wild-type in response to shade treatment, while it did not change in the *shadow-1* mutant. The *GhD2* gene has been reported to accelerate leaf senescence in rice in response to drought and dark treatment (Liu *et al*. 2016; Fan *et al*. 2022). The *NTRC* gene was down-regulated in the wild-type in response to shade treatment, while it did not change in the *shadow-1* mutant. The *NTRC* gene plays a positive role in shade tolerance in plants, as ntrc mutants in Arabidopsis showed more susceptibility to dark and exhibited leaf senescence when grown under short-day conditions (Lepisto *et al*. 2008; Pérez-Ruiz *et al*. 2006; Noroozipoor *et al*. 2020). Additionally, we observed an up-regulation of the *PLS2* gene in the wild-type but a down-regulation in the *shadow-1* mutant under shade conditions. The *PLS2* gene is known to be essential for leaf senescence in plants (Wang *et al*. 2018; Gong *et al*. 2019). The *VPE3* gene plays a crucial role in programmed cell death and leaf senescence in plants under the stress (van Baarlen *et al*. 2007; Yamada *et al*. 2009; Zhang *et al*. 2020; Xiao *et al*. 2021). In our study, the elevation of *VPE3* gene expression was observed in the wild-type under the shade conditions, while it was down-regulated in the *shadow-1* mutant. Suppression of the *VPE3* gene in rice enhances salt tolerance, resulted in more chlorophyll retention and reduced senescence compared to wild-type (Lu *et al*. 2016). GA up-regulates the *VPE3* gene expression level in rice (Xiao *et al*. 2021). *VPE3* gene mediates GA-induced programmed cell death in rice, suggesting a cross-talk between *VPE3* and GA (Zhang *et al*. 2020). Under the shade conditions, the GA1 level was increased in wild-type and decreased in the *shadow-1* mutant, which might change *VPE3* gene expression levels as well. Overall, the differential expression of these senescence-related genes supports the idea that the *shadow-1* mutant has an altered senescence pathway compared to wild-type, which may contribute to its shade tolerance. Additionally, the observed changes in GA, CKs, and ET hormone levels in the mutants compared to the wild-type suggest that these hormones may play important roles in regulating senescence-related gene expression in plants.

## 5. Conclusion

In conclusion, our study provides valuable insights into the molecular basis underlying shade tolerance and tillering in turfgrass. We have demonstrated the possible roles of plant hormones CKs, GAs, ET, and ABA, as well as the tillering inhibitor genes *MAX2*, *CRY1*, and *SnRK1*, under shade conditions in both wild-type and dwarf mutant plants.

## Author Contributions

RK, HY and WL performed shade experiments under field and greenhouse conditions. RK performed RNAseq data analysis and qPCR experiments. GY and XJ performed hormone analysis. RK wrote the original draft of this manuscript. RK, HY, YL and HD reviewed and edited the manuscript. YL supervised the experiments and data interpretations and manuscript writing.

## Acknowledgments

This project is financially supported by the Storrs Agricultural Station and the USDA Hatch Program to YL.

## References

Al-Mana F A, Al-Yafrsi M A. 2021. Tolerance of some warm-season turfgrasses to compaction under shade and sunlight conditions in Riyadh, Saudi Arabia. Saudi Journal of Biological Sciences, 28 (1), pp.1133–1140.

Ashikari M, Sakakibara H, Lin S, Yamamoto T, Takashi T, Nishimura A, Angeles E R, Qian Q, Kitano H, Matsuoka M. 2005. Cytokinin oxidase regulates rice grain production. Science, 309(5735), pp.741–745.

Baena-González E, Lunn J E. 2020. SnRK1 and trehalose 6-phosphate–two ancient pathways converge to regulate plant metabolism and growth. Current Opinion in Plant Biology, 55, pp.52–59.

Bahmani I, Hazard L, Varlet-Grancher C, Betin M, Lemaire G, Matthew C, Thom ER. 2000. Differences in tillering of long-and short-leaved perennial ryegrass genetic lines under full light and shade treatments. Crop Science, 40(4), pp.1095–1102.

Barbier F, Péron T, Lecerf M, Perez-Garcia M D, Barrière Q, Rolčík J, Boutet-Mercey S, Citerne S, Lemoine R, Porcheron B, Roman H. 2015. Sucrose is an early modulator of the key hormonal mechanisms controlling bud outgrowth in Rosa hybrida. Journal of experimental botany, 66(9), pp.2569–2582.

Beall F D, Yeung E C, Pharis R P. 1996. Far-red light stimulates internode elongation, cell division, cell elongation, and gibberellin levels in bean. Canadian Journal of Botany, 74(5), pp.743–752.

Belda-Palazon B, Adamo M, Valerio C, Ferreira L J, Confraria A, Reis-Barata D, Rodrigues A, Meyer C, Rodriguez P L, Baena-González E. 2020. A dual function of SnRK2 kinases in the regulation of SnRK1 and plant growth. Nature Plants, 6(11), pp.1345–1353.

Blakeslee J J, Peer W A, Murphy A S. 2005. Auxin transport. Current opinion in plant biology, 8(5), pp.494–500.

Bou-Torrent J, Galstyan A, Gallemí M, Cifuentes-Esquivel N, Molina-Contreras M J, Salla-Martret M, Jikumaru Y, Yamaguchi S, Kamiya Y, Martínez-García J F. 2014. Plant proximity perception dynamically modulates hormone levels and sensitivity in Arabidopsis. Journal of experimental botany, 65(11), pp.2937–2947.

Brewer P B, Dun E A, Ferguson B J, Rameau C, Beveridge C A. 2009. Strigolactone acts downstream of auxin to regulate bud outgrowth in pea and Arabidopsis. Plant physiology, 150(1), pp.482–493.

Byrne S L, Nagy I, Pfeifer M, Armstead I, Swain S, Studer B, Mayer K, Campbell J D, Czaban A, Hentrup S, Panitz, F. 2015. A synteny based draft genome sequence of the forage grass Lolium perenne. The Plant Journal, 84(4), pp.816–826.

Carrow R N. 1980. Influence of Soil Compaction on Three Turfgrass Species 1. Agronomy Journal, 72(6), pp.1038–1042.

Cheng J, Hill C B, Shabala S, Li C, Zhou M. 2022. Manipulating GA-related genes for cereal crop improvement. International journal of molecular sciences, 23(22), p.14046.

Chugh V, Mishra V, Sharma V, Kumar M, Ghorbel M, Kumar H, Rai A, Kumar R. 2024. Deciphering Physio-Biochemical Basis of Tolerance Mechanism for Sesame (Sesamum indicum L.) Genotypes under Waterlogging Stress at Early Vegetative Stage. Plants, 13(4), p.501.

Cline M G. 1994. The role of hormones in apical dominance. New approaches to an old problem in plant development. Physiologia plantarum, 90(1), pp.230–237.

Das D St, Onge K R, Voesenek L A, Pierik R, Sasidharan R. 2016. Ethylene-and shade-induced hypocotyl elongation share transcriptome patterns and functional regulators. Plant Physiology, 172(2), pp.718–733.

Fan X, Liu J, Zhang Z, Xi Y, Li S, Xiong L, Xing Y. 2022. A long transcript mutant of the rubisco activase gene RCA upregulated by the transcription factor Ghd2 enhances drought tolerance in rice. The Plant Journal, 110(3), pp.673–687.

Ferguson B J, Beveridge C A. 2009. Roles for auxin, cytokinin, and strigolactone in regulating shoot branching. Plant physiology, 149(4), pp.1929–1944.

Fichtner F, Barbier F F, Feil R, Watanabe M, Annunziata M G, Chabikwa T G, Höfgen R, Stitt M, Beveridge C A, Lunn J E. 2017. Trehalose 6 phosphate is involved in triggering axillary bud outgrowth in garden pea (Pisum sativum L.). The Plant Journal, 92(4), pp.611–623.

Filipe O, De Vleesschauwer D, Haeck A, Demeestere K, Höfte M. 2018. The energy sensor OsSnRK1a confers broad-spectrum disease resistance in rice. Scientific Reports, 8(1), p.3864.

Franklin K A, Whitelam G C. 2005. Phytochromes and shade-avoidance responses in plants. Annals of botany, 96(2), pp.169–175.

Gan S, Amasino R M. 1995. Inhibition of leaf senescence by autoregulated production of cytokinin. science, 270(5244), pp.1986–1988.

Ganie S A, Molla K A, Henry R J, Bhat K V, Mondal T K. 2019. Advances in understanding salt tolerance in rice. Theoretical and Applied Genetics, 132, pp.851–870.

Gardner D S, Taylor J A. 2002. Change over time in quality and cover of various turfgrass species and cultivars maintained in shade. HortTechnology, 12(3), pp.465–469.

Glanz-Idan N, Lach M, Tarkowski P, Vrobel O, Wolf S. 2022. Delayed leaf senescence by upregulation of cytokinin biosynthesis specifically in tomato roots. Frontiers in Plant Science, 13, p.922106.

Gong P, Luo Y, Huang F, Chen Y, Zhao C, Wu X, Li K, Yang X, Cheng F, Xiang X, Wu C. 2019. Disruption of a Upf1-like helicase-encoding gene OsPLS2 triggers light-dependent premature leaf senescence in rice. Plant molecular biology, 100, pp.133–149.

Grinberg N F, Lovatt A, Hegarty M, Lovatt A, Skøt K P, Kelly R, Blackmore T, Thorogood D, King R D, Armstead I, Powell W. 2016. Implementation of genomic prediction in Lolium perenne (L.) breeding populations. Frontiers in plant science, 7, p.133.

Gu R, Fu J, Guo S, Duan F, Wang Z, Mi G, Yuan L. 2010. Comparative expression and phylogenetic analysis of maize cytokinin dehydrogenase/oxidase (CKX) gene family. Journal of Plant Growth Regulation, 29, pp.428–440.

Hedden P. 2020. The current status of research on gibberellin biosynthesis. Plant and Cell Physiology, 61(11), pp.1832–1849.

Hisamatsu T, King R W, Helliwell C A, Koshioka M. 2005. The involvement of gibberellin 20-oxidase genes in phytochrome-regulated petiole elongation of Arabidopsis. Plant physiology, 138(2), pp.1106–1116.

Ishikawa S, Maekawa M, Arite T, Onishi K, Takamure I, Kyozuka J. 2005. Suppression of tiller bud activity in tillering dwarf mutants of rice. Plant and Cell Physiology, 46(1), pp.79–86.

Jiang Y, Huang B. 2001. Drought and heat stress injury to two cool season turfgrasses in relation to antioxidant metabolism and lipid peroxidation. Crop science, 41(2), pp.436–442.

Johnson X, Brcich T, Dun E A, Goussot M, Haurogné K, Beveridge C A, Rameau C. 2006. Branching genes are conserved across species. Genes controlling a novel signal in pea are coregulated by other long-distance signals. Plant physiology, 142(3), pp.1014–1026.

Kebrom T H, Brutnell T P, Finlayson, S A, 2010. Suppression of sorghum axillary bud outgrowth by shade, phyB and defoliation signalling pathways. *Plant*, Cell & Environment, 33(1), pp.48–58.

Kebrom T H, Brutnell T P. 2007. The molecular analysis of the shade avoidance syndrome in the grasses has begun. Journal of experimental botany, 58(12), pp.3079–3089.

Kebrom T H, Mullet J E. 2016. Transcriptome profiling of tiller buds provides new insights into PhyB regulation of tillering and indeterminate growth in sorghum. Plant physiology, 170(4), pp.2232–2250.

Kim J, Park S J, Lee I H, Chu H, Penfold C A, Kim J H, Buchanan-Wollaston V, Nam H G, Woo H R, Lim P O. 2018. Comparative transcriptome analysis in Arabidopsis ein2/ore3 and ahk3/ore12 mutants during dark-induced leaf senescence. Journal of Experimental Botany, 69(12), pp.3023–3036.

Kudo T, Kiba T, Sakakibara H. 2010. Metabolism and long distance translocation of cytokinins. Journal of integrative plant biology, 52(1), pp.53–60.

Kumar H, Chugh V, Kumar M, Gupta V, Prasad S, Kumar S, Singh C M, Kumar R, Singh B K, Panwar G, Kumar M. 2023a. Investigating the impact of terminal heat stress on contrasting wheat cultivars: a comprehensive analysis of phenological, physiological, and biochemical traits. Frontiers in Plant Science, 14, p.1189005.

Kumar R, Chanda B, Adkins S, Kousik C S. 2024. Comparative transcriptome analysis of resistant and susceptible watermelon genotypes reveals the role of RNAi, callose, proteinase, and cell wall in squash vein yellowing virus resistance. Frontiers in Plant Science, 15, p.1426647.

Kumar R, Kamuda T, Budhathoki R, Tang D, Yer H, Zhao Y, Li Y. 2022. Agrobacterium-and a single Cas9-sgRNA transcript system-mediated high efficiency gene editing in perennial ryegrass. Frontiers in Genome Editing, 4, p.960414.

Kumar R, Sahi G K, Kaur R, Khanna R, Choudhary O P, Mangat G S, Singh K. 2013. Tolerance response of wild and cultivated Oryza species under iron deficiency condition. Crop Improvement, 40(2), pp.168–172.

Kumar R, Yer H. 2023. Genetic Variation in Tolerance to Iron Deficiency among Species of Oryza. Crops, 3(3), pp.184–194.

Kumar S, Kumar H, Gupta V, Kumar A, Singh C M, Kumar M, Singh A K, Panwar G S, Kumar S, Singh A K, Kumar R. 2023b. Capturing agro-morphological variability for tolerance to terminal heat and combined heat–drought stress in landraces and elite cultivar collection of wheat. Frontiers in Plant Science, 14, p.1136455.

Kurepin L V, Emery R N, Pharis R P, Reid D M. 2007. Uncoupling light quality from light irradiance effects in Helianthus annuus shoots: putative roles for plant hormones in leaf and internode growth. Journal of experimental botany, 58(8), pp.2145–2157.

Le Moigne M A, Guérin V, Furet P M, Billard V, Lebrec A, Spíchal L, Roman H, Citerne S, Morvan-Bertrand A, Limami A, Vian A. 2018. Asparagine and sugars are both required to sustain secondary axis elongation after bud outgrowth in Rosa hybrida. Journal of plant physiology, 222, pp.17–27.

Leduc N, Roman H, Barbier F, Péron T, Huché-Thélier L, Lothier J, Demotes-Mainard S, Sakr, S., 2014. Light signaling in bud outgrowth and branching in plants. Plants, 3(2), pp.223–250.

Lepistö A, Kangasjärvi S, Luomala E M, Hännikäinen K, Brader G, Rintamäki E. 2008. Chloroplastic NADPH thioredoxin reductase mediates photoperiod-dependent development of leaves in Arabidopsis. In Photosynthesis. Energy from the Sun: 14th International Congress on Photosynthesis (pp. 1303–1306). Springer Netherlands.

Li J, Xu P, Zhang B, Song Y, Wen S, Bai Y, Ji L, Lai Y, He G, Zhang D. 2023. Paclobutrazol promotes root development of difficult-to-root plants by coordinating auxin and abscisic acid signaling pathways in Phoebe bournei. International Journal of Molecular Sciences, 24(4), p.3753.

Li W, Katin-Grazzini L, Gu X, Wang X, El-Tanbouly R, Yer H, Thammina C, Inguagiato J, Guillard K, McAvoy R J, Wegrzyn, J, 2017. Transcriptome analysis reveals differential gene expression and a possible role of gibberellins in a shade-tolerant mutant of perennial ryegrass. Frontiers in Plant Science, 8, p.868.

Li W, Katin-Grazzini L, Krishnan S, Thammina C, El-Tanbouly R, Yer H, Merewitz E, Guillard K, Inguagiato J, McAvoy R J, Liu Z. 2016. A novel two-step method for screening shade tolerant mutant plants via dwarfism. Frontiers in Plant Science, 7, p.1495.

Lin M T, Occhialini A, Andralojc P J, Parry M A, Hanson M R. 2014. A faster Rubisco with potential to increase photosynthesis in crops. Nature, 513(7519), pp.547–550.

Lin Q, Wu F, Sheng P, Zhang Z, Zhang X, Guo X, Wang J, Cheng Z, Wang J, Wang H Wan J. 2015. The SnRK2-APC/CTE regulatory module mediates the antagonistic action of gibberellic acid and abscisic acid pathways. Nature Communications, 6(1), p.7981.

Liu D, Tang D, Xie M, Zhang J, Zhai L, Mao J, Luo C, Lipzen A, Zhang Y, Savage E, Yuan G. 2023. Agave REVEILLE1 regulates the onset and release of seasonal dormancy in Populus. Plant physiology, 191(3), pp.1492–1504.

Liu J, Nie B, Yu B, Xu F, Zhang Q, Wang Y, Xu W. 2023. Rice ubiquitin conjugating enzyme OsUbc13 negatively regulates immunity against pathogens by enhancing the activity of OsSnRK1a. Plant Biotechnology Journal, 21(8), pp.1590–1610.

Liu J, Shen J, Xu Y, Li X, Xiao J, Xiong L. 2016. Ghd2, a CONSTANS-like gene, confers drought sensitivity through regulation of senescence in rice. Journal of experimental botany, 67(19), pp.5785–5798.

Ljung K, Bhalerao R P, Sandberg G. 2001. Sites and homeostatic control of auxin biosynthesis in Arabidopsis during vegetative growth. The plant journal, 28(4), pp.465–474.

Lu W, Deng M, Guo F, Wang M, Zeng Z, Han N, Yang Y, Zhu M, Bian H. 2016. Suppression of OsVPE3 enhances salt tolerance by attenuating vacuole rupture during programmed cell death and affects stomata development in rice. Rice, 9, pp.1–13.

Lunn J E, Feil R, Hendriks J H, Gibon Y, Morcuende R, Osuna D, Scheible W R, Carillo P, Hajirezaei M R, Stitt M. 2006. Sugar-induced increases in trehalose 6-phosphate are correlated with redox activation of ADPglucose pyrophosphorylase and higher rates of starch synthesis in Arabidopsis thaliana. Biochemical Journal, 397(1), pp.139–148.

Lush W M. 1990. Turf growth and performance evaluation based on turf biomass and tiller density. Agronomy Journal, 82(3), pp.505–511.

Lyu X, Cheng Q, Qin C, Li Y, Xu X, Ji R, Mu R, Li H, Zhao T, Liu J, Zhou Y. 2021. GmCRY1s modulate gibberellin metabolism to regulate soybean shade avoidance in response to reduced blue light. Molecular plant, 14(2), pp.298–314.

Mao X, Li Y, Rehman S U, Miao L, Zhang Y, Chen X, Yu C, Wang J, Li C, Jing R. 2020. The sucrose non-fermenting 1-related protein kinase 2 (SnRK2) genes are multifaceted players in plant growth, development and response to environmental stimuli. Plant and Cell Physiology, 61(2), pp.225–242.

McSteen P. 2009. Hormonal regulation of branching in grasses. Plant Physiology, 149(1), pp.46–55.

Miyawaki K, Matsumoto Kitano M, Kakimoto T. 2004. Expression of cytokinin biosynthetic isopentenyltransferase genes in Arabidopsis: tissue specificity and regulation by auxin, cytokinin, and nitrate. The plant journal, 37(1), pp.128–138.

Müller D, Leyser O. 2011. Auxin, cytokinin and the control of shoot branching. Annals of botany, 107(7), pp.1203–1212.

Nordström A, Tarkowski P, Tarkowska D, Norbaek R, Åstot C, Dolezal K, Sandberg G. 2004. Auxin regulation of cytokinin biosynthesis in Arabidopsis thaliana: a factor of potential importance for auxin–cytokinin-regulated development. Proceedings of the National Academy of Sciences, 101(21), pp.8039–8044.

Noroozipoor A, Aghdasi M, Sadeghipour H R. 2020. Differential carbohydrate dynamics in Arabidopsis wild-type and ntrc mutant after trehalose feeding. Acta physiologiae plantarum, 42, pp.1–13.

Ongaro V, Leyser O. 2008. Hormonal control of shoot branching. Journal of experimental botany, 59(1), pp.67–74.

Panigrahy M, Ranga A, Das J, Panigrahi K C. 2019. Shade tolerance in Swarnaprabha rice is associated with higher rate of panicle emergence and positively regulated by genes of ethylene and cytokinin pathway. Scientific Reports, 9(1), p.6817.

Parr T W. 1981, July. A population study of a sports turf system. In Proceedings 4th Int. Turfgrass Res. Conf., Ontario Agricultural College, Guelph, Canada (pp. 143–150).

Pearce S. 2021. Towards the replacement of wheat ‘Green Revolution’genes. Journal of Experimental Botany, 72(2), pp.157–160.

Perez-Ruiz J M, Spínola M C, Kirchsteiger K, Moreno J, Sahrawy M, Cejudo F J. 2006. Rice NTRC is a high-efficiency redox system for chloroplast protection against oxidative damage. The Plant Cell, 18(9), pp.2356–2368.

Petersen K, Kolmos E, Folling M, Salchert K, Storgaard M, Jensen C S, Didion T, Nielsen K K. 2006. Two MADS box genes from perennial ryegrass are regulated by vernalization and involved in the floral transition. Physiologia Plantarum, 126(2), pp.268–278.

Pierik R, Cuppens M L, Voesenek L A, Visser E J. 2004a. Interactions between ethylene and gibberellins in phytochrome-mediated shade avoidance responses in tobacco. Plant Physiology, 136(2), pp.2928–2936.

Pierik R, Whitelam G C, Voesenek L A, De Kroon H, Visser E J. 2004b. Canopy studies on ethylene insensitive tobacco identify ethylene as a novel element in blue light and plant– plant signalling. The Plant Journal, 38(2), pp.310–319.

Rameau C, Bertheloot J, Leduc N, Andrieu B, Foucher F, Sakr S. 2014. Multiple pathways regulate shoot branching. Frontiers in plant science, 5, p.741.

Rameau C. 2010. Strigolactones, a novel class of plant hormone controlling shoot branching. Comptes rendus biologies, 333(4), pp.344–349.

Rana K, Yuan J, Liao H, Banga S S, Kumar R, Qian W, Ding Y. 2022. Host-induced gene silencing reveals the role of Sclerotinia sclerotiorum oxaloacetate acetylhydrolase gene in fungal oxalic acid accumulation and virulence. Microbiological Research, 258, p.126981.

Robson P R, Donnison I S, Wang K, Frame B, Pegg S E, Thomas A, Thomas H. 2004. Leaf senescence is delayed in maize expressing the Agrobacterium IPT gene under the control of a novel maize senescence enhanced promoter. Plant Biotechnology Journal, 2(2), pp.101–112.

Roman H, Girault T, Barbier F, Péron T, Brouard N, Pěnčík A, Novák O, Vian A, Sakr S, Lothier J, Le Gourrierec J. 2016. Cytokinins are initial targets of light in the control of bud outgrowth. Plant Physiology, 172(1), pp.489–509.

Sachs T, Thimann K V. 1967. The role of auxins and cytokinins in the release of buds from dominance. American Journal of Botany, 54(1), pp.136–144.

Schäfer M, Brütting C, Meza-Canales I D, Großkinsky D K, Vankova R, Baldwin I T, Meldau S. 2015. The role of cis-zeatin-type cytokinins in plant growth regulation and mediating responses to environmental interactions. Journal of experimental botany, 66(16), pp.4873–4884.

Shang Q, Wang Y, Tang H, Sui N, Zhang X, Wang F. 2021. Genetic, hormonal, and environmental control of tillering in wheat. The Crop Journal, 9(5), pp.986–991.

Shen H, Luong P, Huq E. 2007. The F-box protein MAX2 functions as a positive regulator of photomorphogenesis in Arabidopsis. Plant physiology, 145(4), pp.1471–1483.

Shildrick J P, Peel C H. 1984. Shoot numbers, biomass and shear strength in smooth-stalked meadow-grass (Poa pratensis). Journal of the Sports Turf Research Institute.

Simon N M, Kusakina J, Fernández-López Á, Chembath A, Belbin F E, Dodd A N. 2018. The energy-signaling hub SnRK1 is important for sucrose-induced hypocotyl elongation. Plant physiology, 176(2), pp.1299–1310.

Stirnberg P, Furner I J, Ottoline Leyser, H M. 2007. MAX2 participates in an SCF complex which acts locally at the node to suppress shoot branching. The Plant Journal, 50(1), pp.80–94.

Stirnberg P, van De Sande K, Leyser H O. 2002. MAX1 and MAX2 control shoot lateral branching in Arabidopsis. Development, 129 (5): 1131–1141

Tang D, Li Y, Zhai L, Li W, Kumar R, Yer H, Duan H, Cheng B, Deng Z, Li Y. 2023. Root predominant overexpression of iaaM and CKX genes promotes root initiation and biomass production in citrus. Plant Cell, Tissue and Organ Culture, 155(1), pp.103–115.

Tsai A Y L, Gazzarrini S. 2014. Trehalose-6-phosphate and SnRK1 kinases in plant development and signaling: the emerging picture. Frontiers in plant science, 5, p.119.

van Baarlen P, Woltering E J, Staats M, van Kan J A. 2007. Histochemical and genetic analysis of host and non host interactions of Arabidopsis with three Botrytis species: an important role for cell death control. Molecular Plant Pathology, 8(1), pp.41–54.

Wang H, Chen W, Eggert K, Charnikhova T, Bouwmeester H, Schweizer P, Hajirezaei M R, Seiler C, Sreenivasulu N, von Wirén N, Kuhlmann M. 2018. Abscisic acid influences tillering by modulation of strigolactones in barley. Journal of Experimental Botany, 69(16), pp.3883–3898.

Wang M, Zhang T, Peng H, Luo S, Tan J, Jiang K, Heng Y, Zhang X, Guo X, Zheng J, Cheng Z. 2018. Rice premature leaf senescence 2, encoding a glycosyltransferase (GT), is involved in leaf senescence. Frontiers in plant science, 9, p.560.

Wang Y, Liu H, Xin Q. 2014. Genome-wide analysis and identification of cytokinin oxidase/dehydrogenase (CKX) gene family in foxtail millet (Setaria italica). The Crop Journal, 2(4), pp.244–254.

Werner T, Köllmer I, Bartrina I, Holst K, Schmülling T. 2006. New insights into the biology of cytokinin degradation. Plant biology, 8(03), pp.371–381.

Werner T, Motyka V, Laucou V, Smets R, Van Onckelen H, Schmu lling T. 2003. Cytokinin-deficient transgenic Arabidopsis plants show multiple developmental alterations indicating opposite functions of cytokinins in the regulation of shoot and root meristem activity. The Plant Cell, 15(11), pp.2532–2550.

Wu K, Xu H, Gao X, Fu X. 2021. New insights into gibberellin signaling in regulating plant growth–metabolic coordination. Current Opinion in Plant Biology, 63, p.102074.

Xiao Y, Zhang L, Zhang H, Feng H, Li Z, Chen H. 2021. Interaction between endogenous H 2 O 2 and OsVPE3 in the GA-induced PCD of rice aleurone layers. Plant Cell Reports, 40, pp.691–705.

Xing J, Xu Y, Tian J, Gianfagna T, Huang B. 2009. Suppression of shade-or heat-induced leaf senescence in creeping bentgrass through transformation with the ipt gene for cytokinin synthesis. Journal of the American Society for Horticultural Science, 134(6), pp.602–609.

Xu Y, Gianfagna T, Huang B. 2010. Proteomic changes associated with expression of a gene (ipt) controlling cytokinin synthesis for improving heat tolerance in a perennial grass species. Journal of Experimental Botany, 61(12), pp.3273–3289.

Yadav U P, Ivakov A, Feil R, Duan G Y, Walther D, Giavalisco P, Piques M, Carillo P, Hubberten H M, Stitt M, Lunn J E. 2014. The sucrose–trehalose 6-phosphate (Tre6P) nexus: specificity and mechanisms of sucrose signalling by Tre6P. Journal of experimental botany, 65(4), pp.1051–1068.

Yamada T, Ichimura K, Kanekatsu M, van Doorn W G. 2009. Homologs of genes associated with programmed cell death in animal cells are differentially expressed during senescence of Ipomoea nil petals. Plant and Cell Physiology, 50(3), pp.610–625.

Yan Y, Zhao N, Tang H, Gong B, Shi Q. 2020. Shoot branching regulation and signaling. Plant Growth Regulation, 92(2), pp.131–140.

Yang C, Li L. 2017. Hormonal regulation in shade avoidance. Frontiers in Plant Science, 8, p.1527.

Yeh S Y, Chen H W, Ng C Y, Lin C Y, Tseng T H, Li W H, Ku M S. 2015. Down-regulation of cytokinin oxidase 2 expression increases tiller number and improves rice yield. Rice, 8, pp.1–13.

Yu J, Qiu H, Liu X, Wang M, Gao Y, Chory J, Tao Y. 2015. Characterization of tub4P287L, a β tubulin mutant, revealed new aspects of microtubule regulation in shade. Journal of integrative plant biology, 57(9), pp.757–769.

Zalewski W, Galuszka P, Gasparis S, Orczyk W, Nadolska-Orczyk A. 2010. Silencing of the HvCKX1 gene decreases the cytokinin oxidase/dehydrogenase level in barley and leads to higher plant productivity. Journal of experimental botany, 61(6), pp.1839–1851.

Zhai H, Xiong L, Li H, Lyu X, Yang G, Zhao T, Liu J, Liu B. 2020. Cryptochrome 1 inhibits shoot branching by repressing the self-activated transciption loop of PIF4 in Arabidopsis. Plant communications, 1(3).

Zhang H, Xiao Y, Deng X, Feng H, Li Z, Zhang L, Chen H. 2020. OsVPE3 mediates GA-induced programmed cell death in rice aleurone layers via interacting with actin microfilaments. Rice, 13, pp.1–16.

Zhang J, Zhang Q, Xing J, Li H, Miao J, Xu B. 2021. Acetic acid mitigated salt stress by alleviating ionic and oxidative damages and regulating hormone metabolism in perennial ryegrass (Lolium perenne L.). Grass Research, 1(1), pp.1–10.

Zhang Y, Wang J, Li Y, Zhang Z, Yang L, Wang M, Zhang Y, Zhang J, Li C, Li L, Reynolds M P. 2023. Wheat TaSnRK2. 10 phosphorylates TaERD15 and TaENO1 and confers drought tolerance when overexpressed in rice. Plant Physiology, 191(2), pp.1344–1364.

Zhuang L, Ge Y, Wang J, Yu J, Yang Z, Huang B. 2019. Gibberellic acid inhibition of tillering in tall fescue involving crosstalks with cytokinins and transcriptional regulation of gene controlling axillary bud outgrowth. Plant Science, 287, p.110168.

